# ERK3 Is Involved in Regulating Cardiac Fibroblast Function

**DOI:** 10.1101/2023.12.05.570171

**Authors:** Pramod Sahadevan, Dharmendra Dingar, Sherin A. Nawaito, Reshma S. Nair, Joëlle Trépanier, Fatiha Sahmi, Yanfen Shi, Marc-Antoine Gillis, Martin G. Sirois, Sylvain Meloche, Jean-Claude Tardif, Bruce G. Allen

**Author notes:** Address correspondence to: Bruce G. Allen Montreal Heart Institute 5000 Belanger St., Montréal, Québec, Canada H1T 1C8.

## Abstract

ERK3/MAPK6, an atypical MAPK, activates MAP kinase-activated protein kinase (MK)-5 in selected cell types. MK5 haplodeficient mice show reduced hypertrophy and attenuated increase in *Col1a1* mRNA in response to increased cardiac afterload. In addition, MK5 deficiency alters cardiac fibroblast function. This study was to determine the effect of reduced ERK3 on cardiac hypertrophy following transverse aortic constriction (TAC) and fibroblast biology. Three wk post-surgery, ERK3, but not ERK4 or p38α, was co-immunoprecipitated with MK5 from both sham and TAC heart lysates. The increase in left ventricular mass and myocyte diameter was lower in TAC-ERK3^+/-^ than TAC-ERK3^+/+^ hearts, whereas ERK3 haploinsufficiency did not alter systolic or diastolic function. Furthermore, the TAC-induced increase in *Col1a1* mRNA abundance was diminished in ERK3^+/-^ hearts. ERK3 immunoreactivity was detected in atrial and ventricular fibroblasts but not myocytes. In both quiescent fibroblasts and ‘activated’ myofibroblasts isolated from adult mouse heart, siRNA-mediated knockdown of ERK3 reduced the TGF-β-induced increase in *Col1a1* mRNA. In addition, intracellular type 1 collagen immunoreactivity was reduced following ERK3 depletion in quiescent fibroblasts but not myofibroblasts. Finally, knocking down ERK3 impaired motility in both atrial and ventricular myofibroblasts. These results suggest that ERK3 plays an important role in multiple aspects of cardiac fibroblast biology.

## 1. Introduction

Multiple factors such as myocardial infarction, aortic stenosis, and hypertension increase the afterload on the heart, which imposes mechanical stress on cardiac walls and triggers a series of responses in the heart, including compensatory hypertrophy. Although the hypertrophic response improves or maintains cardiac function in the short-term, if the precipitating condition persists, in the long-term, the myocardium undergoes a series of maladaptive responses that may lead to heart failure (1, 2). Although various signal transduction pathways such as the mitogen-activated protein kinase (MAPK) pathways, calcineurin-nuclear factor of activated T cells (NFAT), and the phosphatidyl inositol 3-kinase (PI3K)/AKT pathway are activated in response to hypertrophic stimuli, the molecular mechanisms causing the deterioration of ventricular function remain unclear (3). In addition to promoting increased cardiomyocyte mass via altering the production of neurohumoral mediators and growth factors, various stimuli may share common molecular, biochemical, and mechanical pathways that detrimentally affect the cardiac structure and function, thus contributing to the development and progression of cardiac remodeling. Another aspect of cardiac remodeling that impairs cardiac function involves cardiac fibroblast activation and subsequent remodeling of the extracellular matrix, resulting in interstitial fibrosis, increased stiffness of the ventricular wall, and impaired diastolic function (4, 5).

The classical MAPKs such as the extracellular signal-regulated kinase 1/2 (ERK1/2), c-Jun N-terminal kinase (JNK) and p38 MAP kinases have been implicated in the development of cardiomyopathies (6, 7). p38 MAP kinases are activated when the heart is exposed to a chronic increase in afterload (8); however, the role of p38 in cardiac disease and repair is complicated. Acute, myocyte-specific expression of a constitutively active MKK3b, leading to activation of endogenous p38, in adult mouse heart resulted in the rapid onset of hypertrophy, interstitial fibrosis, contractile dysfunction, and mortality within 8 days (9). Alternatively, fibroblast-targeted deletion of p38α resulted in 100% mortality following myocardial infarction (10). Targets of p38 include the MAP kinase-activated protein kinases (MKs)−2, −3, and −5. Globally knocking out MK2 delays but does not prevent or reduce hypertrophy and does not alter the extent of interstitial fibrosis at up to 8 weeks after mice are exposed to increased afterload resulting from constriction of the transverse aorta (TAC) (11), whereas it prevents or delays the onset of diabetic cardiomyopathy (12). In contrast, cardiac hypertrophy is attenuated in MK5 haploinsufficient (MK5^+/-^) mice 8 weeks post-TAC as is the increase in *Col1a1* levels within the ventricular myocardium (13). Scar size is reduced in male MK5^+/-^ mice following myocardial infarction (14). Finally, reduced MK5 expression increases collagen secretion, reduces proliferation, and impairs the motility and contraction of cardiac ventricular fibroblasts (14, 15).

Atypical MAPKs such as extracellular-regulated kinase (ERK)3/MAPK6 and ERK4/MAPK4 have become of interest for their role in regulating various cellular activities including cell proliferation, migration, and metabolism. Although MK5 was originally shown to be activated by p38, it has also been shown to be activated by both ERK3 and ERK4 (16-18). ERK3 and ERK4 are expressed in the heart; however, their roles in determining cardiac structure, function or remodeling remain unclear (17, 19). ERK3, but not ERK4 or p38α, co-immunoprecipitate from mouse heart lysates with MK5: the presence of ERK3 or absence of p38α immunoreactivity in MK5 immunoprecipitates is unaffected by increased afterload, where p38 is activated (19). The present study to examine the effect of reduced ERK3 expression on cardiac hypertrophy in response to a chronic increase in afterload induced by constriction of the transverse aorta was undertaken using ERK3 haplodeficient (ERK3^+/-^) mice. Hypertrophy was attenuated as was the increase in *Col1a1* mRNA. Motility was reduced in cardiac myofibroblasts following small interfering (si)RNA-mediated knockdown of ERK3 as was the ability of TGF-β to increase *Col1a1* mRNA. Hence, ERK3 plays an important role in regulating cardiac fibroblast function.

## 2. Materials and methods

### 2.1. Materials

Medium 199 and Fungizone were from Sigma-Aldrich, fetal bovine serum (FBS) and trypsin were from Gibco Laboratories (Life Technologies, Inc.), and penicillin-streptomycin solution was from Multicell. All standard culture plates were from Sarstedt, Inc. Collagen-coated hydrogel-bound (8-kPa) polystyrene 35 mm (#PS35-COL-8) and 6-well (#SW6-COL-8) plates were from Matrigen. PCR Primers were from Invitrogen. SDS-polyacrylamide gel electrophoresis reagents, nitrocellulose, and Bradford Protein Assay reagents were from Bio-Rad Laboratories. Anti-ERK3 antibodies were from Abcam (#ab53277). Anti-GAPDH antibodies were from ThermoFisher Scientific (#AM4300). Rabbit anti-mouse type 1 collagen antibodies (#203002) were from MD Bioproducts Inc. Horseradish peroxidase (HRP)-conjugated secondary antibodies were from Jackson ImmunoResearch Laboratories. Collagenase type 1 (#LS004196) was from Worthington Biochemical Corporation. All other reagents were of analytical grade or best grade available. Type 1 (≥18.2 MΩ/cm) water was used throughout these studies.

### 2.2. ERK3 Haplodeficient Mice

The generation of ERK3 knock-out (ERK3^-/-^) mice has been described previously (20). Heterozygous ERK3^+/-^ mice are fertile and show no overt phenotype. Twelve-week-old male ERK3^+/+^ and ERK3^+/-^ littermates were used for mouse studies. All animal experiments were approved by the local ethics committee and performed according to the guidelines of the Canadian Council on Animal Care.

### 2.3. Isolation and Culture of Cardiac Ventricular Fibroblasts

Cardiac ventricular fibroblasts were isolated from 12–14-week-old C57BL/6 mice as described earlier (21, 22). Briefly, mice were sacrificed by cervical dislocation. The heart was excised and placed in sterile PBS (137 mM NaCl, 2.7 mM KCl, 4.2 mM Na_2_HPO_4_·H_2_O, 1.8 mM KH_2_PO_4_, pH 7.4) at 37 °C. After removal of the atria, ventricular tissue was macerated using scissors and subjected to a series of digestions in dissociation medium (116.4 mM NaCl, 23.4 mM HEPES, 0.94 mM NaH_2_PO_4_·H_2_O, 5.4 mM KCl, 5.5 mM dextrose, 0.4 mM MgSO_4_, 0.1 mM CaCl_2_, 1 mg/ml BSA, 0.5 mg/ml collagenase type IA, 1 mg/ml trypsin, and 0.020 mg/ml pancreatin, pH 7.4). Digestion was at 37 °C on an orbital shaker. Digests were centrifuged at 1500 rpm for 5 min and the pellet was re-suspended in 4 ml of M199 media supplemented with 10% fetal bovine serum (FBS) and 2% penicillin/streptomycin. The cell suspension was plated onto two ‘complaint’ 35 mm collagen-coated soft (8-kPa) hydrogel-bound polystyrene plates or ‘non-compliant’ 35 mm uncoated standard culture dishes (stiffness >10^7^-kPa) to obtain fibroblasts or activated fibroblasts (myofibroblasts), respectively, and incubated in a humidified incubator at 37 °C in a 5% CO_2_ atmosphere for 150 minutes. The medium was changed after 150 min to remove unattached cells and debris and every 48 h thereafter. Cultures from passage 2 were used when they reached 80% confluency. Cells were then washed with PBS and starved with serum-free M199 media for 24 hours prior to treatment with 1 μM Ang II, 1 μM TGF-β, or 10% serum in M199 media.

### 2.4. RNAi depletion of ERK3

Fibroblasts from C57BL/6 mice were depleted of ERK3 using small interfering RNA (siRNA), referred to herein as ERK3-kd. These experiments were performed using mice with the same genetic background. Cells were seeded into 12-well plates at 8 x 10^4^ cells/well. Twenty-four hour post sub-culture, the media was replaced with Opti-MEM containing Ambion Silencer-Select ERK3 siRNA (5 pmol, catalog number: 4390771 ID: S78451) and Lipofectamine 2000 (2µl, Invitrogen) and cells incubated for 19 h. Following a media change, cells were incubated in M199 media containing 10% FBS for 12 hours prior to initiating an experiment.

### 2.5. Scratch Wound Assays

After siRNA transfection, fibroblasts were incubated in serum-free M199 media for 12 h and processed for scratch assays. Briefly, an artificial wound was created in each well by scratching the cell monolayer using a 100 μl pipette tip. Wells were then carefully washed with PBS to remove detached cells and cell debris. The medium was then replaced with 1 ml of fresh serum-free M199 medium alone for control wells or M199 medium containing 1 μM Ang II, 1 μM TGF-β, or 1% serum. Images at time zero and at 24 h later were captured using a Nikon inverted phase-contrast microscope with a digital camera (4x magnification). The area of the wound was quantified using ImageJ software (version 1.49p) and the reduction in wound area was calculated using the following formula: relative wound closure (in %) = 100 – ([wound area at 24 h/original wound area] × 100).

### 2.6 Transverse Aortic Constriction

For baseline assessments of cardiac structure and function, 12-week-old mice underwent transthoracic echocardiography (see below). The following day, transverse aortic constriction (TAC) was done as previously described (11, 13). Briefly, mice were sedated with isoflurane gas plus buprenorphine (0.05 mg/kg, intraperitoneal injection) and animal body temperature was maintained between 36 °C and 37 °C throughout the procedure. A non-absorbable 7.0LJsilk suture was used to constrict the transverse aorta between the innominate and left common carotid arteries, around a 27-gauge custom-made stainless needle. Once the suture was tied, the needle was removed. Sham animals underwent the same surgical procedure but without the placement of a ligature. The surgeon who performed the surgery was blinded as to the genetic status of the mice. Three weeks post-surgery, transthoracic echocardiography was repeated to analyze cardiac structure and function. On the following day, cardiac function was assessed using a Millar Micro-Tip pressure catheter. Mice were then sacrificed, the hearts removed, heart weighed, snap-frozen in liquid nitrogen-chilled 2-methyl butane, and stored at −80 °C.

### 2.7. Echocardiographic Imaging

Transthoracic echocardiography was performed using a 10-14-MHz i13 probe and the Vivid 7 Ultrasound machine (GE Healthcare Ultrasound, Horten, Norway). Mice were lightly anesthetized with isoflurane. Left ventricular structure and function were assessed as described previously (11, 13).

### 2.8. Histological Analyses

Hearts were embedded in Tissue-Tek O.C.T. compound (Sakura Finetek USA, Inc.) and transverse cryosections (8 μm) of the ventricles were prepared and stained with Masson’s trichrome. Images were taken at 10X. Collagen content was quantified using Image Pro Premier Software version 9.2 (Media Cybernetics) and expressed as a percentage of the surface area. Perivascular collagen was excluded from the measurements. To determine cardiac myocyte size, the diameter of the myocytes was assessed in trichrome-stained cryosections using Image Pro Plus.

### 2.9. RNA Extraction and Quantitative PCR

Total cellular RNA was extracted from cultured cardiac fibroblasts, myofibroblasts and Transverse cryosections (14 μm) of murine ventricular myocardium using RNeasy Micro kits (Qiagen Inc.) with minor modifications. Total RNA was isolated by vortexing tissue section in 300 μl TRIzol reagent (Sigma) and 66.7 pg/μl carrier RNA and for cardiac fibroblasts 1 ml TRIzol reagent per well of 12 well plate for 30 s. After incubating at ambient temperature for 5min, 60 μl and 200 μl chloroform for tissue section and cardiac fibroblasts, respectively, were added, and samples were again vortexed and maintained at ambient temperature for an additional 2-3 min. After centrifugation for 15 min at 18,300 xg and 4 °C, the upper aqueous phase was collected, diluted with an equal volume of 70% ethanol, and total RNA purified on Qiagen columns according to the manufacturer’s instructions. Finally total RNA was eluted with 14 μl and 20 μl of distilled, RNAse-free water, respectively, for tissue sections and cardiac fibroblasts. As the quantity of RNA recovered was too small to permit quantification of total RNA, cDNA synthesis was performed with 11 μl of the isolated total RNA from tissue sections and 300 ng of total RNA from cardiac fibroblasts in 20 μl reaction volume as described previously (19). The primers used are shown in Table 1.

**Table 1.**
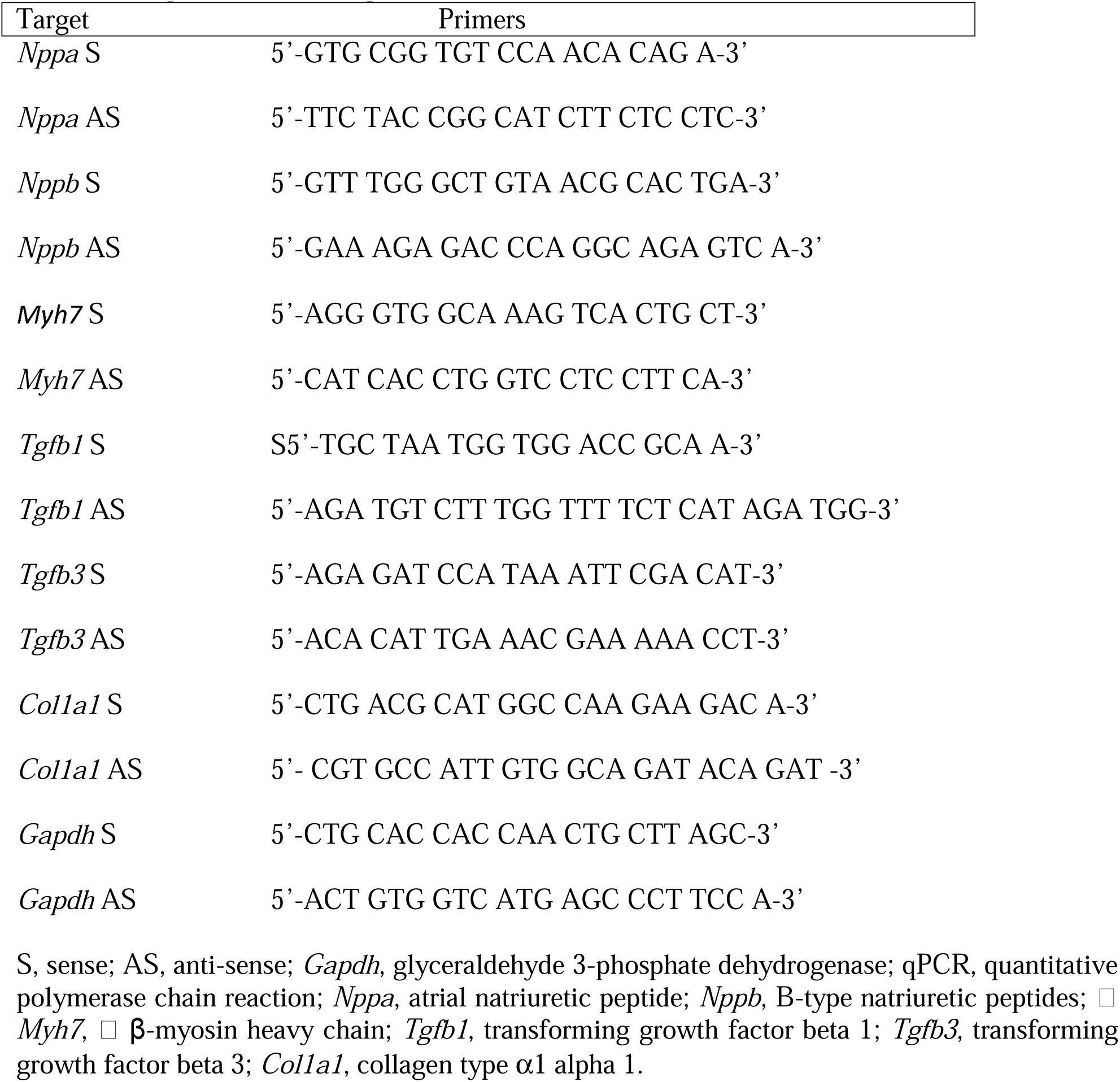
The primers used for qPCR.

### 2.10. Precipitation of Proteins from Conditioned Media

Spend media was collected from cultured fibroblasts and myofibroblasts and 1 ml of media was precipitated by adding 10 μg of bovine serum albumin (BSA) and then adjusting to 10% (w/v) trichloroacetic acid (TCA) using a 100% (w/v) TCA stock solution. The acidified medium was incubated on ice for 1 h and then centrifuged at 15,000 rpm at 4 °C for 30 min. The supernatant was carefully aspirated, and the pellet was resuspended in 30 μl of 1X SDS-PAGE sample buffer. Tris base (3 μl of a 1.0 M stock) was added to neutralize the pH and the samples were heated at 70 °C for 90 s, cooled, centrifuged at room temperature for 1 minute at 14,000 rpm, and loaded onto 6% acrylamide SDS-PAGE gels.

### 2.11. Zymography

The enzymatic activity of MMP-2 and MMP-9 present in the medium was performed with gelatine zymography analyses. Spend media was collected from cultured fibroblasts and 1 ml of media was precipitated by adding 10 μg of bovine serum albumin (BSA) and then adjusting to 10% (w/v) trichloroacetic acid (TCA) using a 100% (w/v) TCA stock solution. The acidified medium was incubated on ice for 1 h and then centrifuged at 15,000 rpm at 4 °C for 30 min. The supernatant was carefully aspirated, and the pellet was resuspended in 30 μl of 1X SDS-PAGE sample buffer without 2-mercaptoethanol and EDTA. Tris base (3 μl of a 1.0 M stock) was added to neutralize the pH. Gelatin zymography was performed under non-reducing conditions on 8% SDS-PAGE copolymerized with 0.2% (w/v) gelatin. Following the electrophoresis, the SDS was removed from the gels by incubating in 2.5% Triton X-100 (three changes, 20 minutes each). Gels were then washed twice with substrate buffer (50 mM Tris-HCl, 150 mM NaCl, 5 mM CaCl_2_, and 0.05% NaN_3_) for 20 minutes each. Zymograms were subsequently incubated in substrate buffer for 24 h at 37 °C. Gels were stained with Coomassie Brilliant Blue and unstained areas corresponding to zones of proteinase activity were visualized after destaining in 2% methanol in 4% acetic acid.

### 2.12. Immunoprecipitation Assays

Co-Immunoprecipitations were performed as described previously (19). Briefly, immediately after the sacrifice of the anesthetized mice, hearts were removed and snap frozen in liquid nitrogen. The heart tissue was pulverized under liquid nitrogen. The tissue powder was resuspended in 1.2 ml of ice-cold lysis buffer (50 mM Tris-Cl (pH 7.5 at 4 °C), 20 mM β-glycerophosphate, 20 mM NaF, 5 mM EDTA, 10 mM EGTA, 1.0% (w/v) TX-100, 1 mM Na_3_VO_4_, 1 μM microcystin LR, 5 mM DTT, 10 μg/ml leupeptin, 0.5 mM PMSF, and 10 mM benzamidine) and homogenized in a 2 ml Potter-Elvehjem tissue grinder (15 passes, on ice). Homogenates were then cleared by centrifugation (100,000 x*g,* 30 min, 4 °C). Prior to immunoprecipitation, A/G^+^-Agarose beads (Santa Cruz Biotechnology) were incubated with 2 μg of anti-MK5 antibody (overnight at 4 °C). Tissue extracts (2 mg) were incubated with antibody-coated beads overnight at 4 °C. Beads were washed thrice with lysis buffer and twice with 50 mM Tris-Cl (pH 7.5 at 4 °C). Beads were then suspended in 20 μl of 2X SDS sample buffer and heated at 70 °C for 90 s and then pelleted by brief centrifugation in a microcentrifuge. The supernatants were immediately loaded onto 10–20% acrylamide-gradient SDS-PAGE gel.

### 2.13. Immunoblot Assays

SDS-PAGE and immunoblotting were performed as described previously (13). Briefly, quiescent fibroblasts and myofibroblasts were lysed in ice-cold LB1 lysis buffer comprising 50 mM Tris-HCl, 20 mM β**-**glycerophosphate, 20 mM NaF, 5 mM EDTA, 10 mM EGTA, 1% TX-100, 1 mM NaVO4, 1 μM microcystin LR, 5 mM DTT, 10 μg/ml leupeptin, 0.5 mM PMSF, and 10 mM benzamidine (pH 7.5 at 4 °C). Samples were separated on SDS-PAGE gels and then transferred onto nitrocellulose membranes. Following the transfer, membranes were rinsed in PBS, fixed with 0.1% (v/v) glutaraldehyde (14), rinsed again with PBS, and blocked in TBST containing 5% non-fat dried milk 5 for 1 h or in 3% bovine serum albumin (BSA) in PBS overnight at 4 °C. Membranes were then incubated with primary antibodies against collagen, ERK3, p-p38, p38α, or GAPDH. Following primary antibody incubation, membranes were washed thrice in TBST and incubated with the appropriate horseradish peroxidase–conjugated secondary antibody. Visualization of immunoreactive bands was achieved by using Western Lightning Plus ECL reagent and Kodak BioMax Light film. Films were digitized using a 2D scanner and band intensities quantified with Image-J software.

### 2.14. Statistical Analysis

The Data are presented as mean ± SEM. For multiple comparisons of means involving a single independent factor such as genotype (ERK3^+/+^, ERK3^+/-^), unpaired Mann-Whitney tests were performed. For multiple comparisons of means involving a combination of 2 independent factors such as surgery (TAC, sham) or genotype (ERK3^+/+^, ERK3^+/-^), 2-way ANOVA followed by Tukey’s post hoc tests were performed. A *P* value < 0.05 was considered significant. Statistical analyses were performed using Prism version 9.5.1 for Mac OS X (GraphPad Software, La Jolla, CA).

## 3. Results

The present study aimed to evaluate the impact of ERK3 haploinsufficiency on pathological cardiac remodeling induced by a chronic pressure overload in mice. Furthermore, this study also compared the effect(s) of reduced ERK3 expression on cardiac fibroblast function.

### 3.1. ERK3 Haplodeficiency Did Not Alter Cardiac Structure and Function

Transthoracic echocardiographic studies were performed to compare 12-wk old male ERK3^+/-^ mice with age-matched male ERK^+/+^ littermates (20). ERK3^+/+^ and ERK3^+/-^ mice showed no significant differences in R-R interval, left ventricular structure, systolic function, peak early transmitral flow velocity (E wave), isovolumetric deceleration time, or myocardial performance index (**Table 2**). Although ERK3^+/-^ and ERK3^+/+^ mice showed no difference in basal lateral or basal septal mitral annular peak systolic velocity (S_L_, S_S_), both basal lateral and basal septal mitral annular peak diastolic velocity was reduced in ERK3^+/-^ mice (E_m_). As a consequence E/E_m_, an indicator of LV filling pressure, increased, suggesting LV compliance may be reduced in ERK3^+/-^ compared to ERK3^+/+^ mice. However, E wave deceleration was unaffected by ERK3 haploinsufficiency.

**Table 2.**
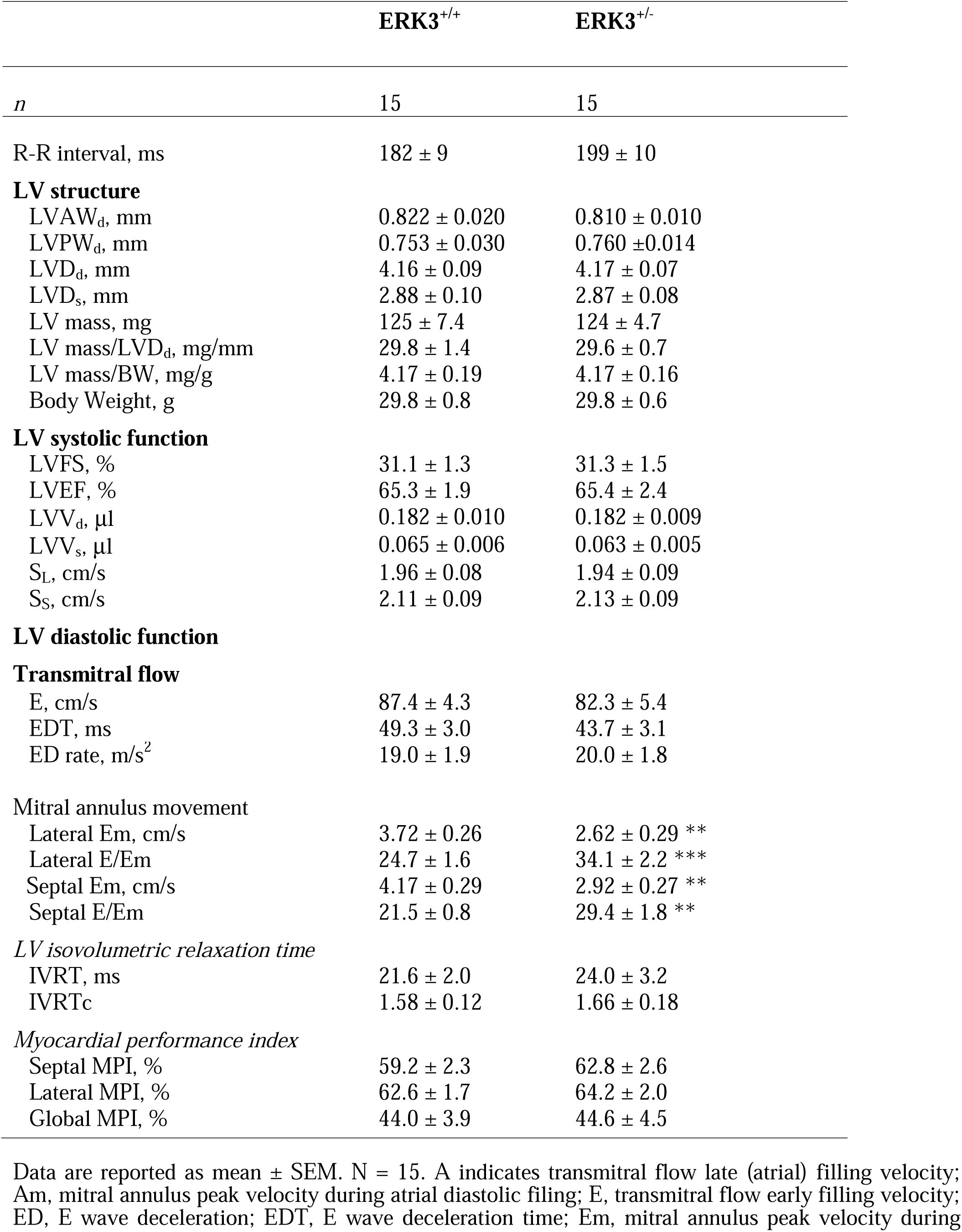
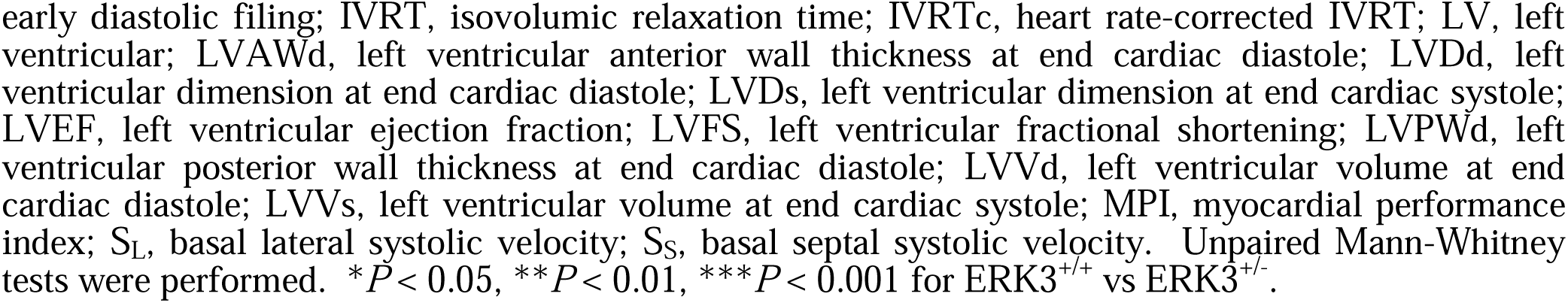
Echocardiographic parameters of 12-week-old ERK3^+/+^ and ERK3^+/-^ mice.

### 3.2. ERK3 Haplodeficiency Attenuated the Hypertrophic Response in Mice Exposed to a Chronic Increase in Afterload

To understand the effect of ERK3 haploinsufficiency on cardiac remodeling in response to increased afterload, 12-week-old male ERK3^+/-^ and ERK3^+/+^ littermate mice (20) underwent surgical constriction of the transverse aorta (TAC) and were sacrificed 3 wk post-surgery. Direct assessment of hemodynamics revealed that systolic blood pressure (BP_max_) as well as peak LV pressure (LVP_max_) were similarly increased in ERK3^+/+^ and ERK3^+/-^ mice following TAC (**Figure 1A**, **Table 3**). Echocardiographic imaging 3-wk post-TAC cross-TAC peak velocity was lower in banded TAC-ERK3^+/-^ mice than TAC ERK3^+/+^ mice (**Table 4**). The ratio of heart weight to tibia length increased in response to TAC; however, ERK3^+/-^ mice did not differ from ERK3^+/+^ mice in either the sham or the TAC groups (**Figure 1B**). The increase in heart weight to tibia length ratio was similar in TAC-ERK3^+/-^ and TAC-ERK3^+/+^ mice. In contrast, the left ventricular anterior wall thickness (LVAW_d_, **Figure 1C**), left ventricular mass (LV mass, **Figure 1D**), ratio of LV mass to left ventricular diameter at diastole (LV mass/LVD_d_, **Figure 1E**), and myocyte diameter (**Figure 1F**) were significantly less in TAC-ERK3^+/-^ mice compared with TAC-ERK3^+/+^ mice. Although the myocardial performance index (MPI) did not differ between ERK3^+/-^ and ERK3^+/+^ mice, 2-way ANOVA indicated a significant interaction between the effects of TAC and genotype on both global and lateral MPI (**Table 4**). Hence, in response to pressure overload, LV hypertrophy was attenuated in ERK3 haplodeficient mice compared to wild-type mice.

**Figure 1.**
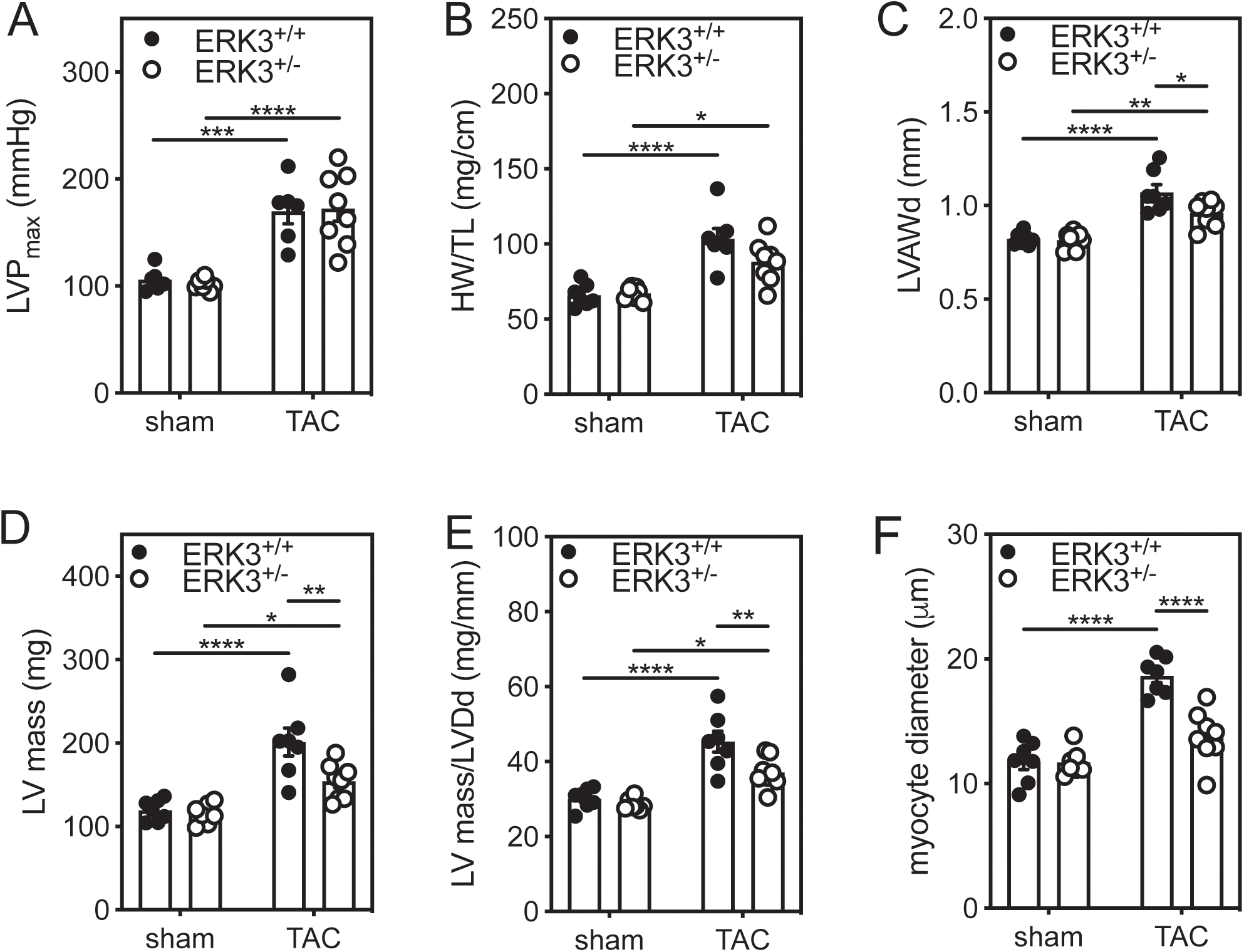
ERK3 Haplodeficiency Attenuates Hypertrophy Induced by Increased Afterload. Cardiac afterload was increased in 12-week-old male ERK3^+/+^ and ERK3^+/-^ littermate mice by constriction of the transverse aorta (TAC). The mice were sacrificed 3 wk post-surgery. Sham-operated animals underwent the identical surgical procedure; however, the aorta was not constricted. (**A**) Left ventricular maximum developed pressure (LVP_max_) was measured using a Millar mikro-tip pressure catheter. (**B**) Heart weight (HW) normalized to tibia length (TL). (**C**) Left ventricular anterior wall thickness in diastole (LVAW_d_), (**D**) left ventricular mass (LV mass), and (**E**) LV mass/left ventricular end-diastolic diameter (LVmass/LVD_d_) were determined by echocardiography (**Table 4**). (**F**) Myocyte diameter was calculated from fixed sections following Masson’s trichrome staining. Shapiro-Wilk tests for normality were performed on all data prior to statistical comparison was by two-way ANOVA, including a factor for surgery (Sham, TAC), a factor for genotype (ERK3^+/+^, ERK3^+/-^), and a surgery x genotype interaction term. The ANOVA was followed by Tukey’s post hoc tests for multiple comparisons. Data from 6-7 mice per group are shown as mean ± S.E.M. \*\*\*\**P* < 0.0001, \*\*\**P* < 0.001, \*\**P* < 0.01, \**P* < 0.05.

**Table 3.** Hemodynamic Parameters of ERK3^+/+^ and ERK3^+/-^ Mice 3 Weeks Post-TAC.

|  | Sham Operation |  | TAC |  |
| --- | --- | --- | --- | --- |
|  | ERK3<sup>+/+</sup> | ERK3<sup>+/-</sup> | ERK3<sup>+/+</sup> | ERK3<sup>+/-</sup> |
| n | 6 | 7 | 6 | 8 |
| BP<sub>max</sub>, mm Hg | 106±5 | 101±1 | 156±11<sup>c</sup> | 155±7<sup>d</sup> |
| LVP<sub>max</sub>, mm Hg | 106±4 | 102±2 | 170±12<sup>c</sup> | 172±12<sup>d</sup> |
| LVP<sub>min</sub>, mm Hg | 3.8±1.6 | 1.3±0.6 | 4.9±1.5 | 3.3±1.3 |
| +dP/dt, mm Hg/s | 6480±362 | 6515±361 | 6969±530 | 8273±616 |
| -dP/dt, mm Hg/s | 5831±447 | 5833±339 | 7148±357 | 8273±672<sup>b</sup> |
| LVEDP, mm Hg | 9.5±1.9 | 6.3±1.0 | 9.9±2.1 | 8.8±1.8 |
| CT, ms | 11.7±0.4 | 14.1±2.1 | 11.1±0.5 | 10.8±0.4 |
| RT, ms | 48.3±2.0 | 48.6±1.3 | 47.4±1.0 | 45.1±1.2 |
| Tau, ms | 9.88±0.46 | 9.08±0.60 | 9.05±0.50 | 8.13±0.56 |
| HR, beats/min | 346±19 | 344±21 | 350±11 | 389±10 |

Data reported as mean ± SEM. Shapiro-Wilk tests for normality were performed. Statistical comparison of means was by two-way ANOVA, including a factor for surgery (sham, TAC), a factor for genotype (ERK3<sup>+/+</sup>, ERK3<sup>+/-</sup>), and a surgery x genotype interaction term. The ANOVA was followed by Tukey's post hoc tests for multiple comparisons. BP<sub>max</sub> indicates maximum systolic blood pressure; CT, contraction time; +dP/dt, maximum rate of left ventricular pressure increase during contraction; -dP/dt, maximum rate of left ventricular decline during relaxation; HR, heart rate; LVEDP, left ventricular end-diastolic pressure; LVP<sub>max</sub>, left ventricular maximum systolic pressure; LVP<sub>min</sub>, left ventricular minimum diastolic pressure; RT, relaxation time; Tau, time constant; and TAC, transverse aortic constriction. <sup>b</sup>*P* < 0.01 vs. sham; <sup>c</sup>*P* < 0.001 vs. sham; <sup>d</sup>*P* < 0.0001 vs. sham.

**Table 4.**
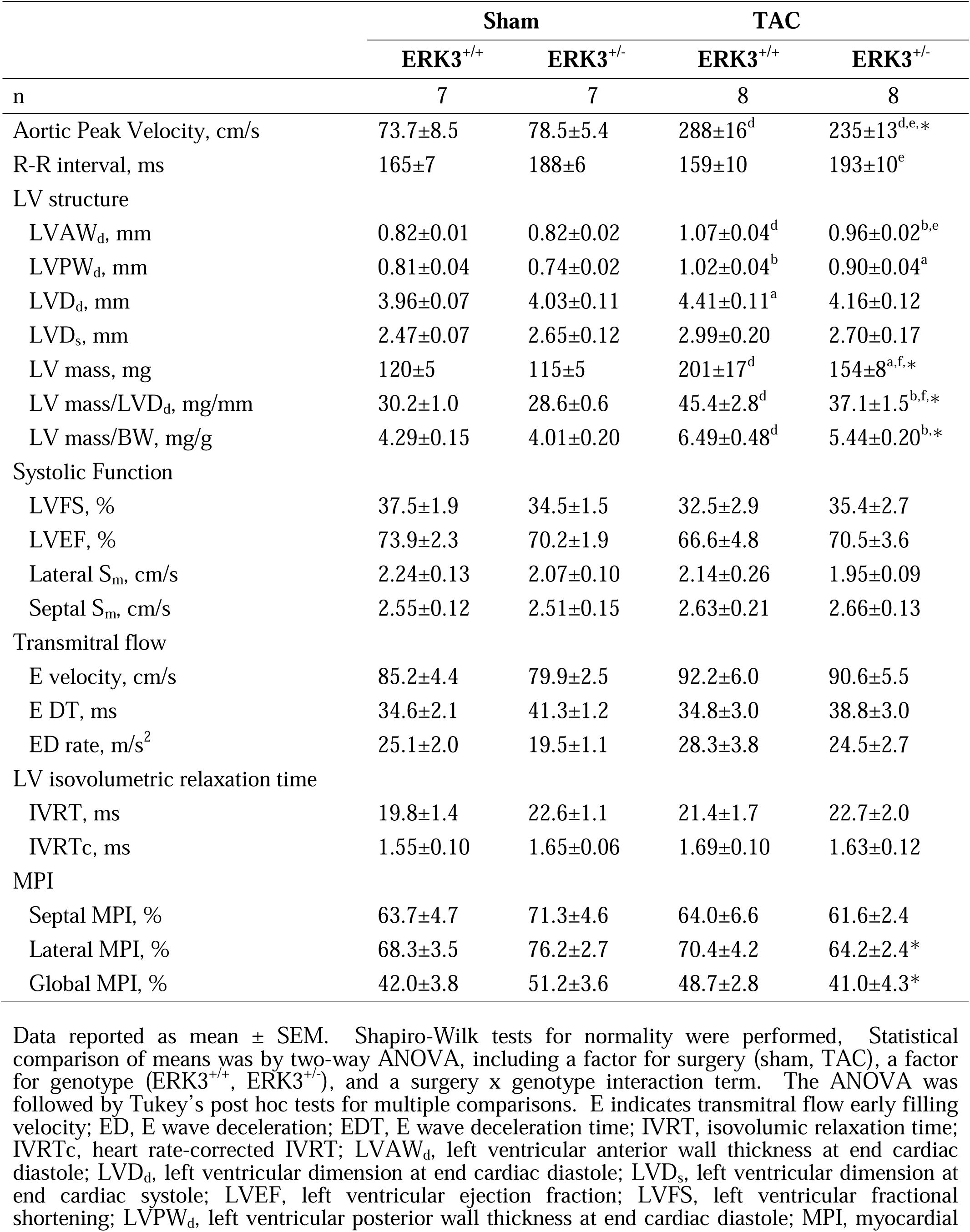
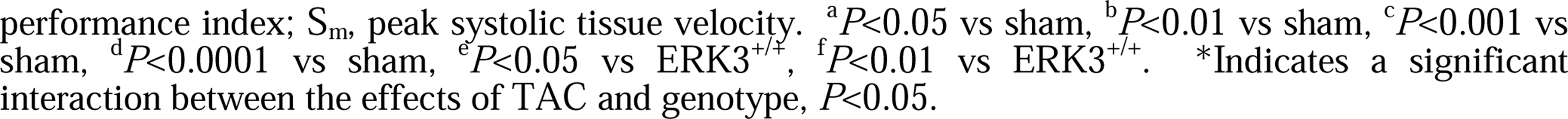
Echocardiographic Parameters of ERK3^+/+^ and ERK3^+/-^ Mice 3 Weeks After TAC.

### 3.3. ERK3 Haplodeficiency Did Not Alter Fetal Gene Re-expression

Reexpression of fetal genes, molecular markers of pathologic cardiac hypertrophy, are commonly found in association with cardiac remodeling. Hence, the abundance of atrial natriuretic peptide (*NppA*), B-type natriuretic peptide (*Nppb*), and β-myosin heavy chain (*Myh7*) mRNA was assessed by qPCR three weeks post-TAC. *Nppb* and *Myh7* mRNAs were more abundant increased significantly in TAC-ERK3^+/+^ and TAC-ERK3^+/-^ hearts, relative to sham mice. However, ERK3 haplodeficiency did not affect the abundance of these transcripts in either sham or TAC mice (**Figure 2A-C**).

**Figure 2.**
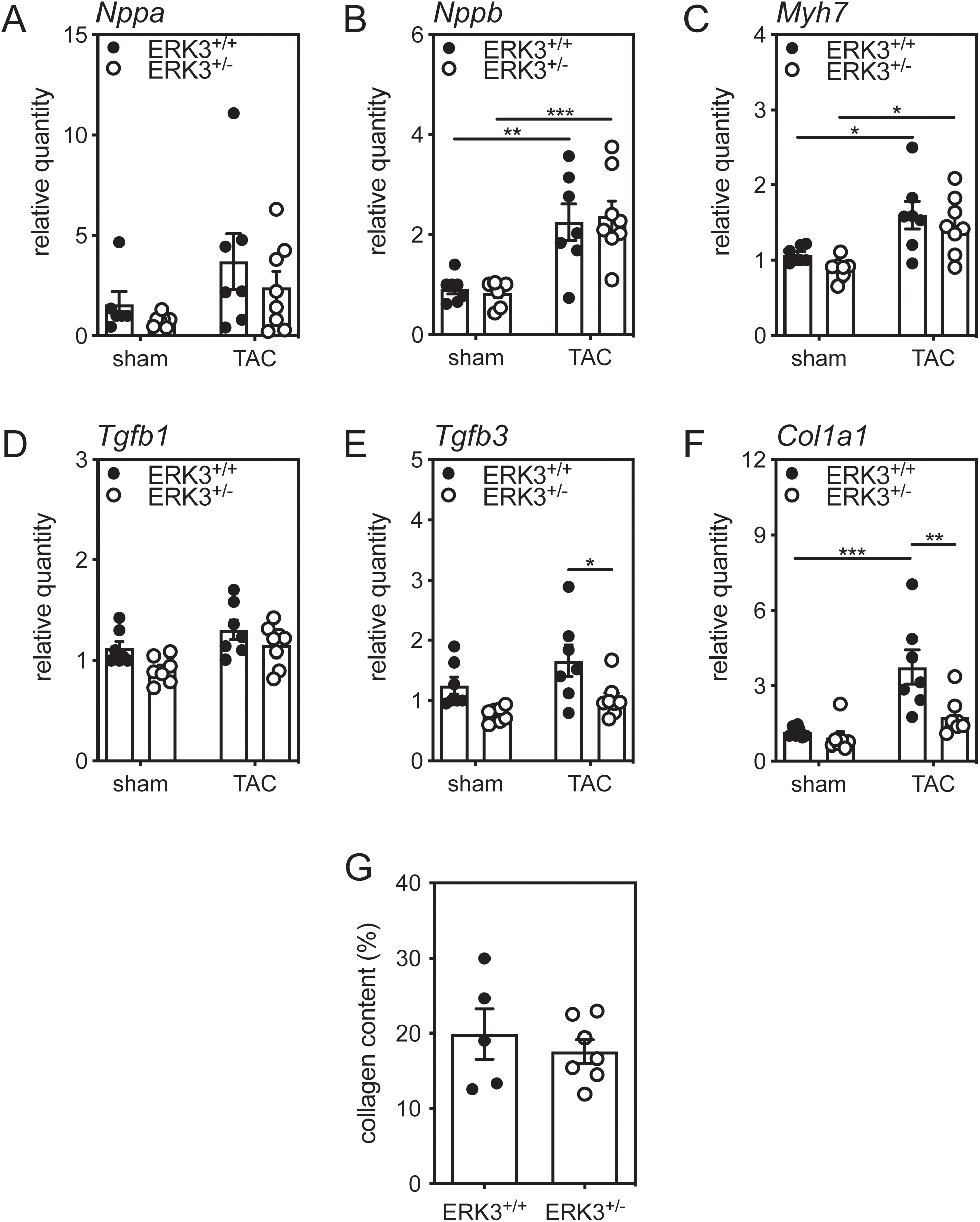
ERK3 Haplodeficiency Does Not Alter Fetal Gene reexpression following TAC but Attenuates the Increase in *Col1a1*. Total RNA was isolated from sham and TAC hearts and the relative abundance of the mRNA for (**A**) A-type natriuretic peptide (*Nppa*), (**B**) B-type natriuretic peptide (*Nppb*), (**C**) β-myosin heavy chain (*Myh7*), (**D**) transforming growth factor-β1 (*Tgfb1*), (**E**) transforming growth factor-β3 (*Tgfb3*), and (**F**) collagen type 1 α1 (*Col1a1*) was quantified by qPCR and normalized to that of glyceraldehyde-3-phosphate dehydrogenase (*Gapdh*). (**G**) Quantitative analysis of the collagen content in the LV wall following Masson’s trichrome staining. For data having 2 independent variables, Shapiro-Wilk tests for normality were performed on all data prior to statistical comparison was by two-way ANOVA, including a factor for surgery (Sham, TAC), a factor for genotype (ERK3^+/+^, ERK3^+/-^), and a surgery x genotype interaction term. The ANOVA was followed by Tukey’s post hoc tests for multiple comparisons. For data sets comprising a single independent variable, statistical analysis was by 2-way unpaired Mann-Whitney test. Data from 6-7 mice per group are shown as mean ± S.E.M. \*\*\**P* < 0.001, \*\**P* < 0.01, \**P* < 0.05.

### 3.4. ERK3 Haplodeficiency Attenuated the TAC-induced increase in *Col1a1* mRNA

We next examined the effect of ERK3 haplodeficiency on markers of fibrosis. The abundance of *Col1a1*, *Tgfb1*, and *Tgfb3* mRNA was assessed by qPCR. No significant differences were observed in the abundance of *Tgfb1* (**Figure 2D**) and that of *Tgfb3* was significantly lower in hearts from TAC-ERK3^+/-^ mice than TAC-ERK3^+/+^ mice whereas sham mice did not differ significantly (**Figure 2E**). Interestingly, the abundance of *Col1a1* mRNA increased in response to TAC in hearts from ERK3^+/+^ mice and this increase was significantly attenuated in hearts from TAC-ERK3^+/-^ mice (**Figure 2F**). Masson’s trichrome staining revealed that the left ventricular myocardial collagen content did not differ significantly between TAC-ERK3^+/-^ and TAC- ERK3^+/+^ mice (Not Shown).

### 3.5. ERK3 and MK5 Co-Immunoprecipitate from Heart 3 Weeks Post-TAC

Although controversial, ERK3, ERK4, and p38α/β MAPK have each been shown to bind to and activate MK5. An earlier study from our group showed that ERK3, but not ERK4 or p38α, co-immunoprecipitated with MK5 from the heart lysates prepared from control, sham-operated, or 3- d post-TAC mouse (19). To ascertain if ERK3-MK5 complexes remain stable or dissociate as cardiac remodeling progresses in response to a chronic increase in afterload, co-immunoprecipitation assays were performed in lysates prepared from sham and TAC hearts 3-wk post-TAC. In each case, ERK3, but not p38α or ERK4 immunoreactivity was detected in the MK5 immunoprecipitates (**Figure 3A**).

**Figure 3.**
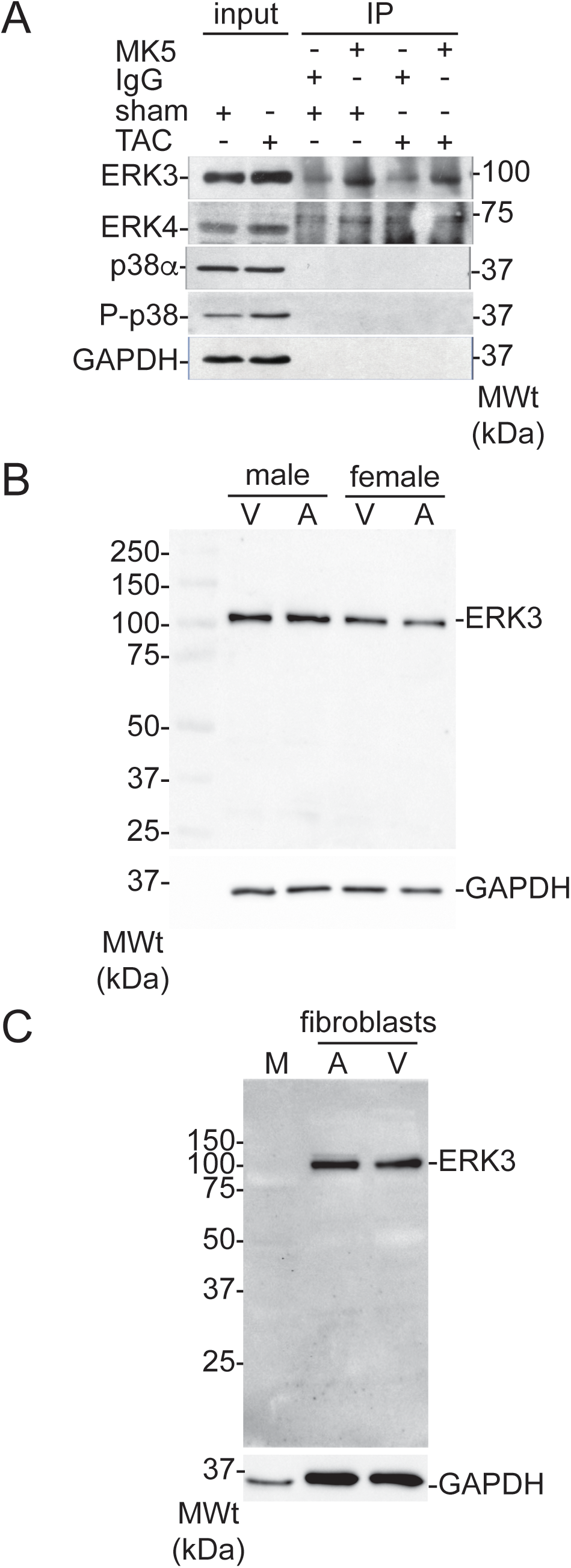
ERK3 Immunoreactivity Co-immunoprecipitates With MK5 in Sham and TAC Hearts and is Detected in Atrial and Ventricular Fibroblasts. (**A**)Mice were sacrificed 3 wk after surgery, the ventricular myocardium isolated, lysates prepared from TAC- or sham-treated hearts and incubated with protein A/G agarose beads loaded with either anti-MK5 antibodies or purified rabbit IgG. After washing, immunoprecipitates were denatured using Laemmli sample buffer, resolved on SDS-PAGE, and transferred onto nitrocellulose membranes. Membranes where then incubated with ERK3-, ERK4-, p38α-, phospho-p38(T180/Y182)-, or GAPDH-specific antibodies. Aliquots (50 μg) of cardiac lysate from TAC and sham hearts were included as controls (input). Results shown are representative of 3 independent experiments using lysates from different hearts for each determination. Numbers on the right indicate the position of the molecular mass markers (in kDa). (**B**) ERK3-immunoreactive bands were detected in the lysates (25 μg) of fibroblasts isolated from the atria (A) and ventricles (V) of wild type male and female mice. The numbers at the left indicate the position or the molecular mass markers (in kDa). GAPDH immunoreactivity was used as a loading control. (**C**) ERK3-immunoreactivity was not detected in the lysates (25 μg) of ventricular myocytes (M) isolated from adult male rat hearts whereas it was detected in atrial (A) and ventricular (V) fibroblasts isolated from adult male rat hearts. The numbers at the left indicate the position or the molecular mass markers (in kDa). GAPDH immunoreactivity was used as a loading control.

### 3.6. Small Interfering (si)RNA-Mediated ERK3 Knockdown Attenuated the ability of TGF-β to increase *Col1a1*

We have shown previously that MK5, the haplodeficiency of which also attenuates both TAC-induced hypertrophy and increased *Col1a1* mRNA, is detected by immunoblotting in adult rat cardiac ventricular fibroblasts but not myocytes (13). Immunoblotting revealed ERK3 immunoreactivity in myofibroblasts of ventricular and atrial origin isolated from adult male and female mice (**Figure 3B**) but not in ventricular myocytes (**Figure 3C**). Hence, we next examined the effect of reduced ERK3 on fibroblast function. We used a RNAi approach to acutely deplete ERK3 (ERK3-kd) in fibroblasts. Immunoblotting revealed that ERK3 siRNA reduced the abundance of ERK3 immunoreactivity by 80-85% (**Figure 4A**) relative to a scrambled RNA (scRNA) control. To study the effects of reduced ERK3 on the phenotype of both quiescent fibroblasts (qFBs) and ‘activated’ myofibroblasts (mFBs), once isolated, fibroblasts were maintained on either compliant (elastic modulus 8-kPa) or rigid, non-compliant (elastic modulus ∼10^7^-kPa) substrates, respectively (22): the abundance of smooth muscle α-actin (αSMA) immunoreactivity, a marker of fibroblast activation, was much greater in fibroblasts maintained on a non-compliant substrate.(**Figure 4B**).

**Figure 4.**
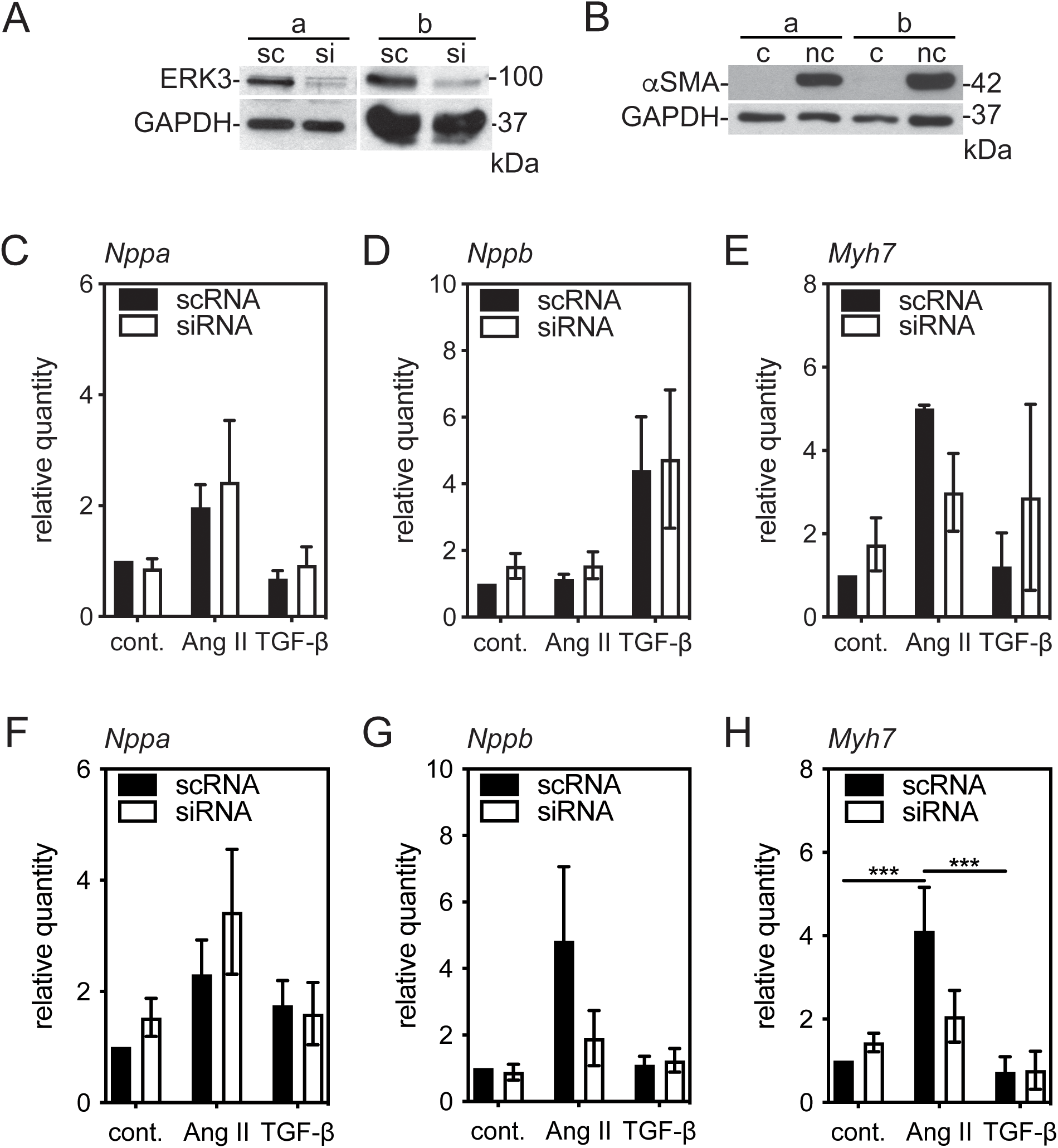
Effect of Small Interfering (si)RNA-Mediated ERK3 Knockdown on the abundance of transcripts for fetal proteins in ventricular fibroblasts and myofibroblasts. Ventricular fibroblasts from ERK3^+/+^ mice were isolated and cultured on ‘compliant’ (c) collagen-coated, hydrogel-bound (elastic modulus 8-kPa) polystyrene plates or ‘non-compliant’ (elastic modulus 10^7^-kPa) uncoated standard cell culture dishes (nc), resulting in quiescent fibroblasts or myofibroblasts, respectively. Quiescent fibroblasts or myofibroblasts were transiently transfected with either siRNA for ERK3 (si) or a scrambled RNA sequence (sc) as described under Methods. In both cases, after 12 h of serum deprivation, cells were incubated in serum-free M199 in the presence or absence of 1 μM angiotensin II (Ang II) or transforming growth factor-β1 (TGF-β) for 24 h. (**A**) Immunoblot assay showing siRNA reduced the abundance of ERK3 immunoreactivity in myofibroblast lysates from two independent experiments using separate cell preparations (a, b). The numbers at the right indicate the position of molecular mass markers (in kDa). (**B**) An immunoblot assay showing the abundance of smooth muscle α-actin (αSMA) immunoreactivity in lysates of fibroblasts that were grown on compliant (c) and non-compliant (nc) substrates. Results from two independent cell preparations (a, b) are shown. The numbers at the right indicate the position of molecular mass markers (in kDa). The relative abundance of the mRNA (**C, F**) A-type natriuretic peptide (*Nppa*), (**D, G**) B-type natriuretic peptide (*Nppb*), and (**E, H**) β-myosin heavy chain (*Myh7*) in quiescent fibroblasts (**C - E**) and myofibroblasts (**F - H**) was evaluated by qPCR and. The data shown was normalized to *Gapdh* mRNA. Data shown are mean ± SEM for n = 3 cell preparations. Statistical analysis was by 2-way ANOVA followed by Tukey’s post hoc tests was performed. ***P < 0.001.

We examined the effect of an ERK3-kd on the abundance of *NppA*, *Nppb*, and *Myh7* mRNA, which increases during pathological hypertrophy in cardiac ventricular qFBs and mFBs. Quiescent fibroblasts and myofibroblasts were transfected with either ERK3-specific siRNA or scRNA and subsequently treated with angiotensin II (Ang II) or TGF-β for 24 h. In qFBs, there was a significant main effect of treatment on the abundance of *Nppa* (F(2, 23) = 4.357, *P* = 0.025) and *Nppb* (F(2, 22) = 7.609, *P* = 0.0031) but not *Myh7* (**Figure 4C-E**). In each case, knocking down ERK3 had no effect. In mFBs, there was a significant main effect of treatment on the abundance of *Nppa* (F(2, 28) = 4.493, *P* = 0.020), *Nppb* (F(2, 29) = 4.785, *P* = 0.016), and *Myh7* (F(2, 22) = 15.10, *P* < 0.0001). In each case, knocking down ERK3 had no main effect; however, there was a significant interaction between the effect of knocking down ERK3 and the effect of treatment on the abundance of Myh7 mRNA (F(2, 22) = 4.836, *P* = 0.0182) in mFBs (**Figure 4F-H**). It should be noted that the abundance of these transcripts in both qFBs and mFBs was very low, with cycle threshold (Ct) values of 35, resulting in highly variable data but indicating that fibroblasts are only, at best, minor contributors to the reexpression of these fetal genes commonly employed as molecular markers of pathological hypertrophy.

As ERK3 haplodeficiency was associated with a reduction in the TAC-induced increase in *Col1a1* mRNA (**Figure 2F**), we examined the effect of knocking down ERK3 on the abundance of *Col1a1* mRNA and type 1 collagen immunoreactivity in qFBs and mFBs. Fibroblasts were transfected with either ERK3-specific siRNA or scRNA, serum starved, and then treated with media alone or supplemented with either 1 μM Ang II or TGF-β. *Col1a1* mRNA, assessed by qPCR, was normalized to that of *Gapdh*. In both qFBs and mFBs, the main effects of both treatment (qFBs, F(2, 17 = 4.819, *P* = 0.022; mFBs, F(2, 18 = 4.830, *P* = 0.021)) and ERK3-kd (qFBs, F(1, 17 = 16.6, *P* = 0.0008; mFBs, F(1, 18 = 6.54, *P* = 0.020)) on the abundance of *Col1a1* mRNA were statistically significant. Knocking down ERK3 prevented TGF-β from increasing the abundance of *Col1a1* mRNA in both qFBs and mFBs (**Figures 5A,B**). A reduction was also observed in qFBs treated with Ang II, which failed to reach significance. The abundance of both intracellular and secreted type 1 collagen immunoreactivity was assessed by immunoblot assay. In qFBs, there was a significant main effect of knocking down ERK3 on the abundance of both intracellular (F(1, 12) = 36.34, *P* = 0.0001) and secreted (F(2, 12)) = 14.78, *P* = 0.0023) type 1 collagen immunoreactivity with a reduction in immunoreactivity being observed in both cases (**Figure 5C,D).** In contrast, there was a significant main effect of treatment on the abundance of secreted (F(1, 12) = 5.678, *P* = 0.0184) but not intracellular collagen immunoreactivity. Furthermore, there was a statistically significant interaction between the effect of ERK3 knockdown and the effect of treatment (Ang II, TGF-β) on the abundance of intracellular type 1 collagen immunoreactivity (F(2, 12) = 11.7, *P* = 0.0016). No effect of treatment or ERK3-kd was observed in mFBs under the conditions employed (**Figures 5E,F**).

**Figure 5.**
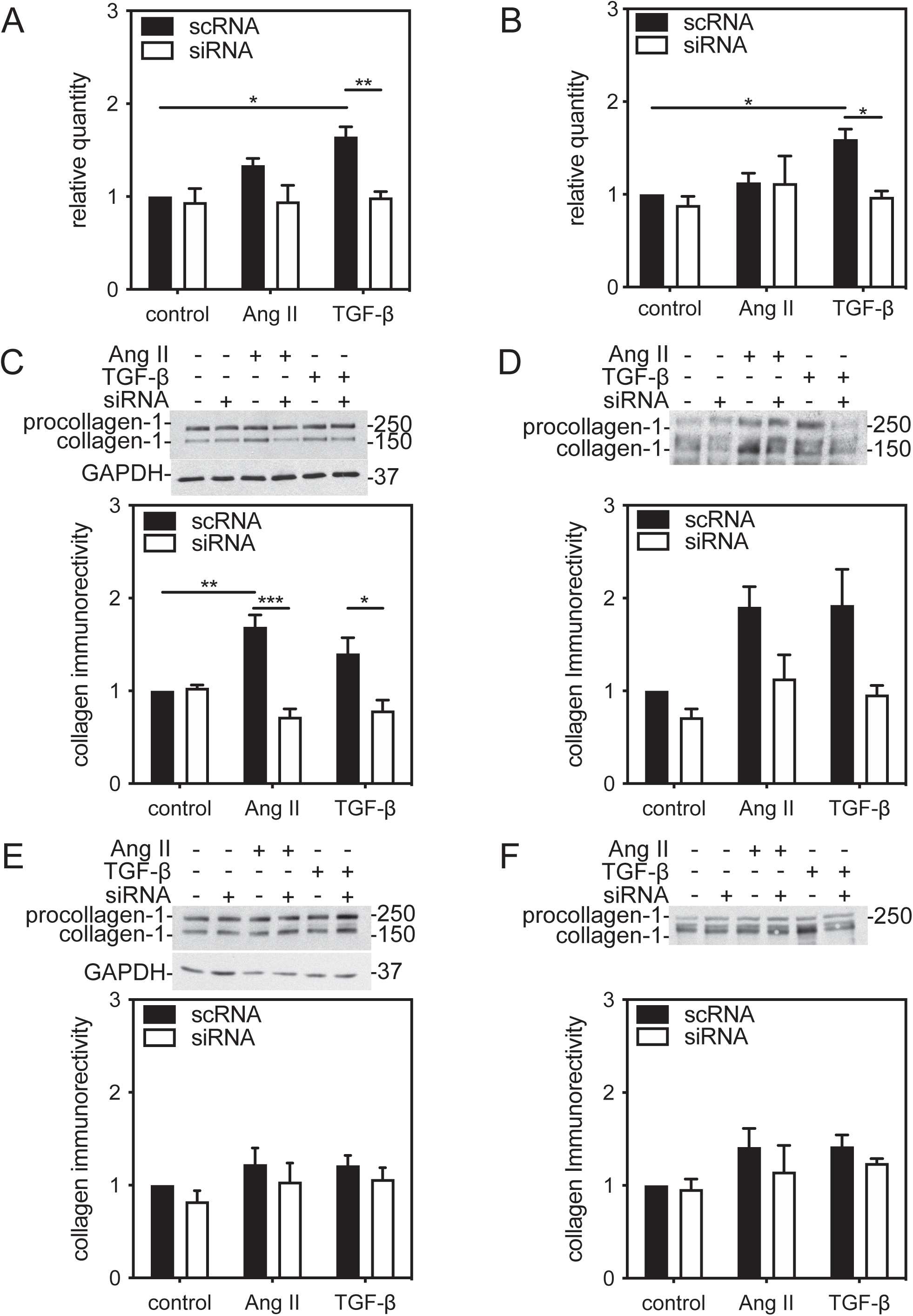
Small interfering (si)RNA-mediated knockdown of ERK3 in fibroblasts reduced the abundance of *Col1a1* mRNA but not type 1 collagen immunoreactivity. Ventricular fibroblasts from ERK3^+/+^ mice were isolated and cultured on (**A, C, D**) ‘compliant’ collagen-coated, hydrogel-bound (elastic modulus 8-kPa) plates or (**B, E, F**) ‘non-compliant’ (elastic modulus 10^7^-kPa) uncoated standard cell culture dishes, resulting in quiescent fibroblasts or myofibroblasts, respectively. Quiescent fibroblasts or myofibroblasts were transiently transfected with either siRNA for ERK3 (siRNA) or a scrambled RNA sequence (scRNA) as described under Methods. In both cases, after 12 h of serum deprivation, cells were incubated in serum-free M199 in the absence (control) or presence of 1 μM angiotensin II (Ang II) or transforming growth factor-β1 (TGF-β) for 24 h. (A, B) *Col1a1* mRNA was quantified by qPCR and normalized *Gapdh* mRNA. (**C-F**) Type 1 collagen immunoreactivity was measured in cell lysates (**C, E**; 20μg) or conditioned media (**D, F**) by immunoblot assay. Results were normalized to cell number and then control fibroblasts transfected with scRNA. Representative immunoblots are shown above the histograms. GAPDH immunoreactivity was employed as a loading control when determining intracellular collagen immunoreactivity. Numbers on the right of the immunoblot images indicate the position of the molecular mass markers (in kDa). Values in the histograms show the mean ± SEM of three independent experiments. Each experiment was performed with fibroblasts isolated from a separate mouse. Statistical analysis was by 2-way ANOVA followed by Tukey’s post hoc tests was performed. \**P* < 0.05, \*\**P* < 0.01, \*\*\**P* < 0.001.

### 3.7. siRNA-Mediated ERK3 Knockdown Had Only Modest Effects on MMP2 and MMP9 Activity

In addition to extracellular matrix components, fibroblasts produce enzymes involved in extracellular matrix maintenance and remodelling (23). To characterize the effects of knocking down ERK3 on MMP activity in cardiac fibroblasts, we examined both cell-associated and secreted MMP2 and MMP9 activity in qFBs and mFBs stimulated with media alone (control) or media supplemented with Ang II or TGF-β by collagen zymogram assay. Although activities corresponding to both latent and mature forms of MMP-2 were present in all groups (**Figures 6,7**), MMP2 activity was highest in the conditioned media from mFBs (**Figure 7C**). In qFBs, there was a significant main effect of treatment (F(2, 12) = 4.08, *P* = 0.044), but not ERK3 knockdown, on cell-associated MMP2 activity. Cell-associated MMP9 activities was unaltered by treatment or by knocking down ERK3 (**Figure 6A,C**) as was secreted MMP9 (**Figure 6D,F**). In contrast, secreted MMP2 activity was reduced in the presence of TGF-β in both scRNA- and siRNA-qFBs (**Figure 6D,E**). Although there was a small, but not significant, reduction in cell-associated MMP2 activity in the presence of Ang II, this was prevented by knocking down ERK3. Furthermore, there was a statistically significant interaction between the effects of ERK3 knockdown and treatment (Ang II, TGF-β) on secreted MMP2 activity (F(2, 18)) = 4.79, *P* = 0.022).

**Figure 6.**
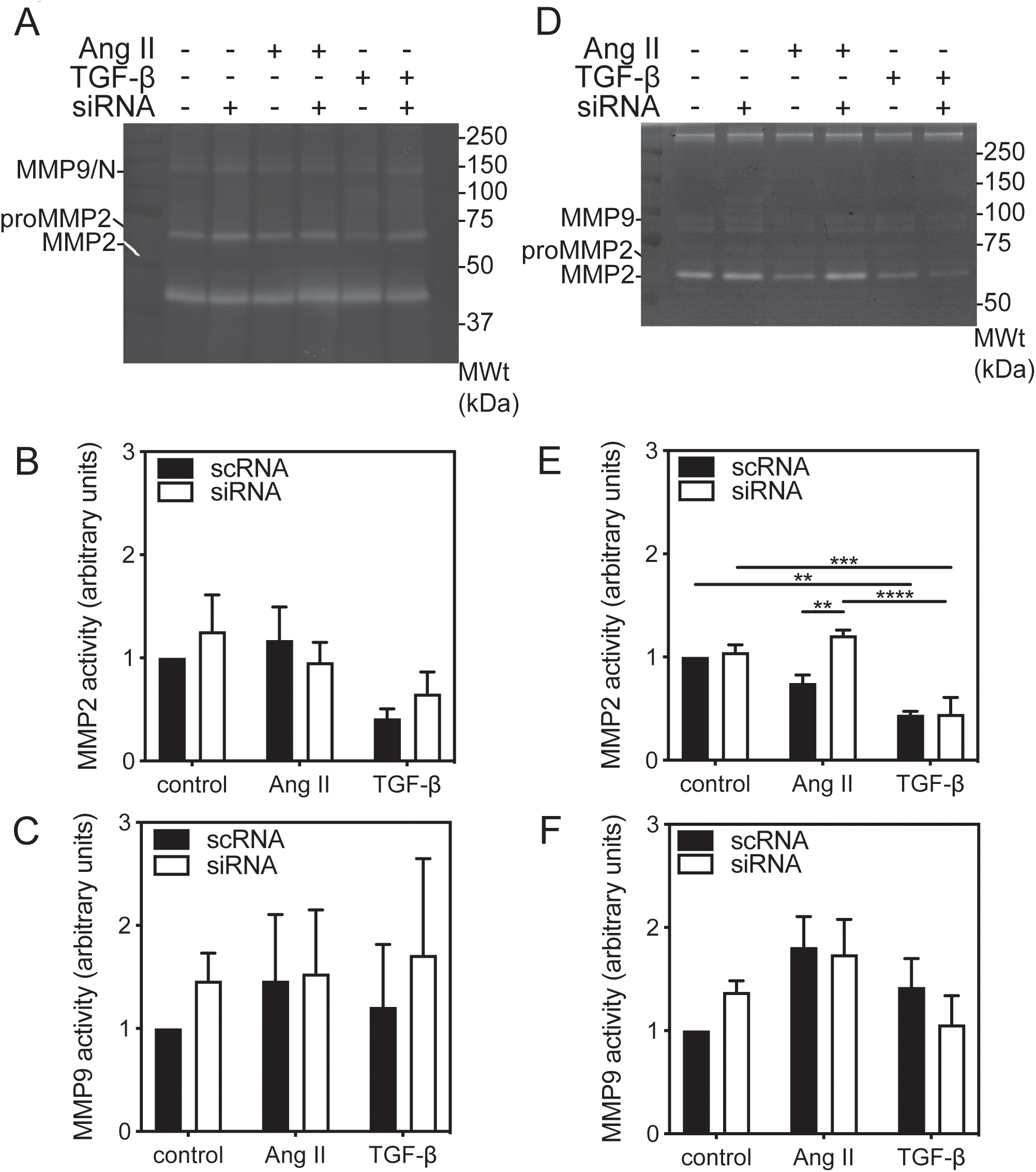
MMP-2 and MMP-9 activity in ERK3-deficient quiescent ventricular fibroblasts. Ventricular fibroblasts isolated from ERK3^+/+^ mice, maintained on ‘compliant’ collagen-coated, hydrogel-bound (elastic modulus 8-kPa) plates, resulting in quiescent fibroblasts, were transiently transfected with either siRNA for ERK3 (siRNA) or a scrambled RNA sequence (scRNA). After 12 h of serum deprivation, cells were incubated in serum-free M199 in the absence (control) or presence of 1 μM angiotensin II (Ang II) or transforming growth factor-β1 (TGF-β) for 24 h, and both the cell lysate (20 μg) and conditioned media used for the measurement of MMP-2 and MMP9 activity by gelatin zymography. (**A**) Representative zymogram showing bands of cell-associated pro-MMP-2, MMP-2, and MMP9-NGAL (MMP9/N) activity. Numbers on the right of the image indicate the position of molecular weight markers (in kDa). Quantitative analysis of MMP-2 activity (**B**) and MMP-9-NGAL activity (**C**) normalized to that of fibroblasts transfected with scRNA and maintained in serum-free media. (**D**) Representative zymogram showing bands of secreted pro-MMP-2, MMP-2, and MMP9 activity. Numbers on the right of the image indicate the position of molecular weight markers (in kDa). Quantitative analysis of MMP-2 activity (**E**) and MMP-9 activity (**F**) normalized to that of fibroblasts transfected with scRNA and maintained in serum-free media. The data shown are mean ± SEM of assessments performed in fibroblasts isolated from 3 different mice. Statistical analysis was by 2-way ANOVA followed Tukey’s post hoc test for multiple comparisons. \*\**P* < 0.01, \*\*\**P* < 0.001, \*\*\*\**P* < 0.0001.

**Figure 7.**
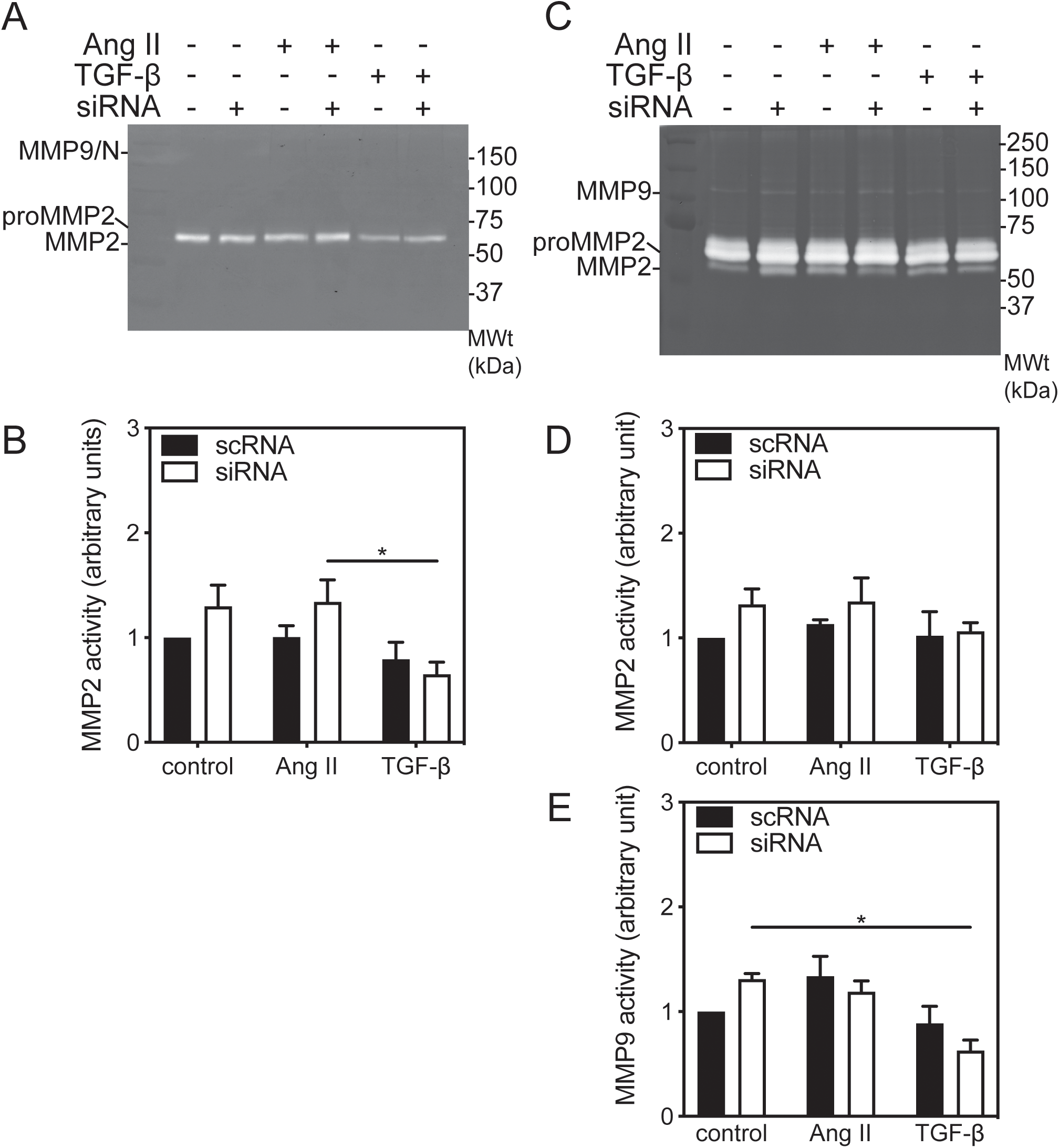
MMP-2 and MMP-9 activity in ERK3-deficient ventricular myofibroblasts. Ventricular myofibroblasts isolated from ERK3^+/+^ mice, maintained on ‘non-compliant’ (elastic modulus 10^7^-kPa) standard cell culture dishes, resulting in myofibroblasts, were transiently transfected with either siRNA for ERK3 (siRNA) or a scrambled RNA sequence (scRNA). After 12 h of serum deprivation, cells were incubated in serum-free M199 in the absence (control) or presence of 1 μM angiotensin II (Ang II) or transforming growth factor-β1 (TGF-β) for 24 h. Both cell lysates (20 μg) and conditioned media used for the measurement of MMP-2 and MMP-9 activity by gelatin zymography. (**A**) Representative zymogram showing bands of cell-associated pro-MMP-2, MMP-2, and MMP-9-NGAL (MMP/N) activity. Numbers on the right of the image indicate the position of molecular weight markers (in kDa). Quantitative analysis of MMP-2 activity (**B**) activity normalized to that of fibroblasts transfected with scRNA and maintained in serum-free media. (**C**) Representative zymogram showing bands of secreted pro-MMP-2, MMP-2, and MMP9 activity. Numbers on the right of the image indicate the position of molecular weight markers (in kDa). Quantitative analysis of MMP-2 activity (**D**) and MMP-9 activity (**E**) normalized to that of fibroblasts transfected with scRNA and maintained in serum-free media. Statistical analysis was by 2-way ANOVA followed Tukey’s post hoc test for multiple comparisons. The data shown are mean ± SEM of assessments performed in fibroblasts isolated from 3 different mice. \**P* < 0.05.

There was a significant main effect of treatment (F(2, 18) = 5.64, *P* = 0.013), but not ERK3 knockdown, on cell-associated MMP2 activity in mFBs. MMP2 activity was unaffected by treatment of scRNA-mFBs with Ang II whereas in both control and Ang II-treated siRNA-mFBs, cell associated MMP2 activity showed a small, but insignificant, increase (**Figure 7A,B**). Interestingly, in siRNA-ERK3 mFBs, cell-associated MMP2 activity was significantly lower when treated with TGF-β than with AngII. No cell-associated MMP9 activity was detected in mFBs(**Figure 7A**). Secreted MMP2 activity was unaffected by treatment or ERK3-knockdown (**Figure 7C,D).** The response observed with secreted MMP9 activity resembled that observed with cell-associated MMP2 activity: there was a significant main effect of treatment (F(2, 12) = 9.83, *P* = 0.003), but not ERK3 knockdown, on secreted MMP9 activity in mFBs. Ang II produced a non-significant increase in secreted MMP9 activity in scRNA-mFBs. In siRNA- mFBs, secreted MMP9 activity was significantly lower when treated with TGF-β than when treated with media alone.

### 3.8. siRNA-Mediated ERK3 Knockdown Reduced Myofibroblast Motility

To further determine the role of ERK3 in fibroblast function, we examined the effect of knocking down ERK3 on the migratory response of the cardiac myofibroblasts, using scratch wound assays, to Ang II, TGF-β, or serum. Ventricular fibroblasts, plated on a non-compliant substrate, were grown to 80% confluence, “wounds” were created, and the cells were incubated for an additional 24 h with or without the addition of angiotensin II, TGF-β or serum to the media. Wound areas were measured at time 0 and after 24 h. After 24 h of culture in serum-free medium, wound closure for cardiac fibroblasts transfected with scRNA or siRNA did not differ significantly (scRNA: 14.7 ± LJ2.3%; siRNA: 17.8 ± 3.3%; n = 7; **Figure 8A,B**). However, after 24 h, wound closure in scRNA-mFBs treated with angiotensin II, TGF-β or serum was significantly greater in comparison with scRNA-mFBs. There was a statistically significant interaction between the effects of treatment (Ang II, TGF-β, serum) and knocking down ERK3 on motility (F(3, 28) = 8.09, *P* = 0.0005).

**Figure 8.**
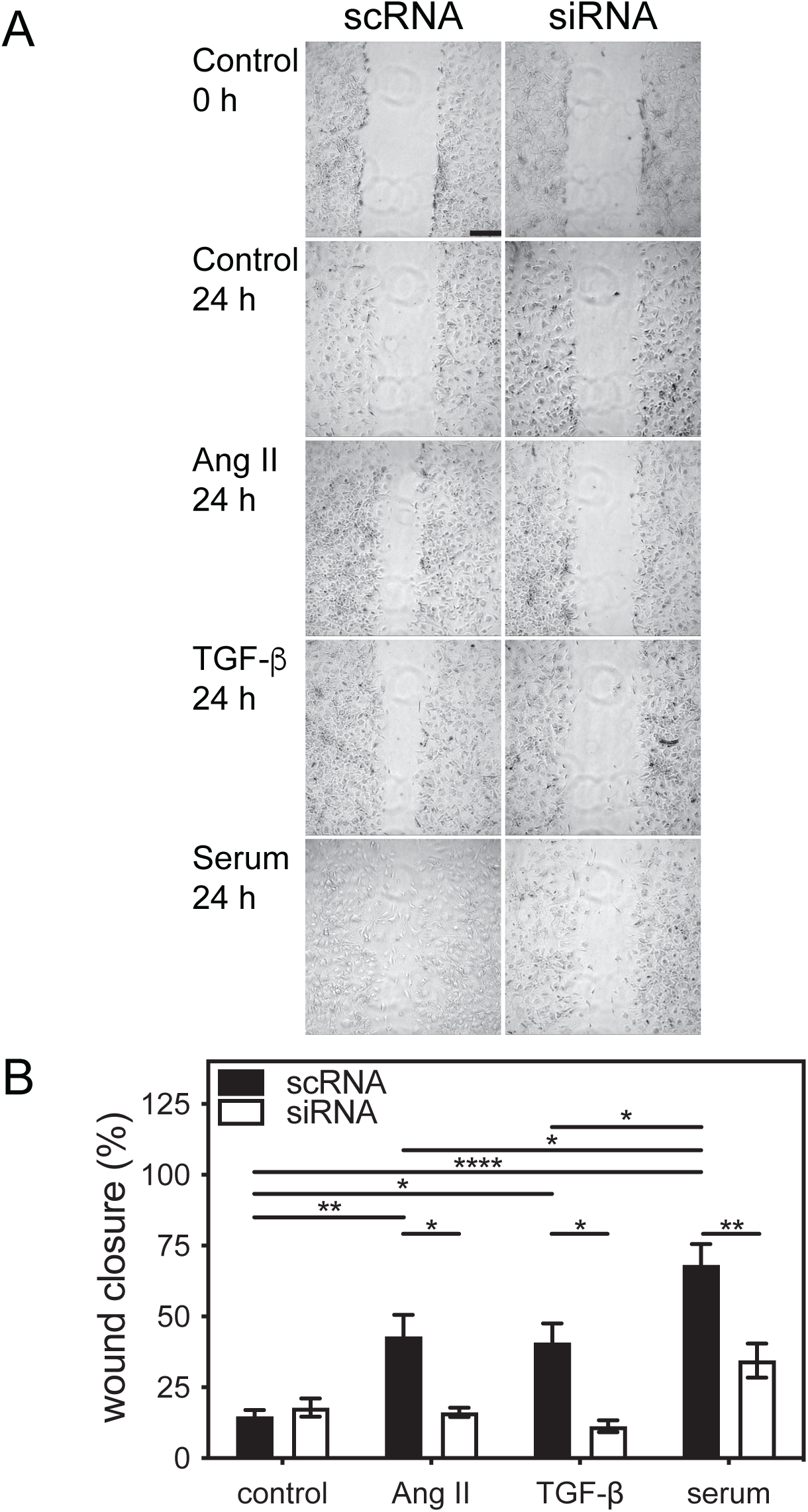
Small interfering (si)RNA-mediated knockdown of ERK3 impairs motility in ventricular myofibroblasts. (**A**) Cardiac fibroblasts were isolated from the ventricular myocardium of wild type mice, split into two, and transfected with either siRNA for ERK3 (si) or a scrambled RNA sequence (sc), and motility was assessed by scratch wound assay. After the scratch was imposed, fibroblasts were cultured in serum-free media alone (control) or supplemented with angiotensin II (Ang II; 1 μM), TGF-β (1 μM), or 1% serum. Open wound areas were measured at times 0 and 24 h, and wound closure was calculated. (**B**) Wound closure after 24 h normalized to the mean closure of the control scRNA group (expressed as %). Statistical analysis was by 2-way ANOVA followed Tukey’s post hoc test for multiple comparisons. The data shown are mean ± SEM of assessments performed in fibroblasts isolated from 3 different mice. Bar, 250 μm. \**P* < 0.05, \*\**P* < 0.01, \*\*\*\**P* < 0.0001.

We next examined the effect of ERK3 knockdown on the motility of fibroblasts isolated from mouse atria. Atrial fibroblasts, plated on a non-compliant substrate, were grown to 80% confluence, “wounds” were created, and the cells were incubated for an additional 24 h with or without the addition of angiotensin II, TGF-β or serum to the media. Wound areas were measured at time 0 and after 24 h. After 24 h of culture in serum-free medium, wound closure for atrial mFBs transfected with scRNA or siRNA did not differ significantly (scRNA: 47.3 ± LJ3.3%; siRNA: 37.8 ± 17.8%; n = 3; **Figure 9A,B**). It was notable that the basal rate of migration for atrial myofibroblasts was markedly greater than ventricular mFBs (**Figure 8**). In addition, although Ang II and serum, but not TGF-β, produced a small increase in atrial mFBs transfected with scRNA, these changes did not reach significance. Knocking down ERK3 in atrial mFBs resulted in significantly lower migration in the presence of Ang II or serum. There was a significant main effect of knocking down ERK3 (F(1, 16) = 25.74, *P* = 0.0001), but not treatment (F(3, 16) = 2.198, *P* = 0.13), on atrial mFB motility. In addition, after 24 h, the cell density outside of the scratch in ERK3-kd atrial mFBs appeared to be lower, suggesting a detrimental effect on either viability or adhesion.

**Figure 9.**
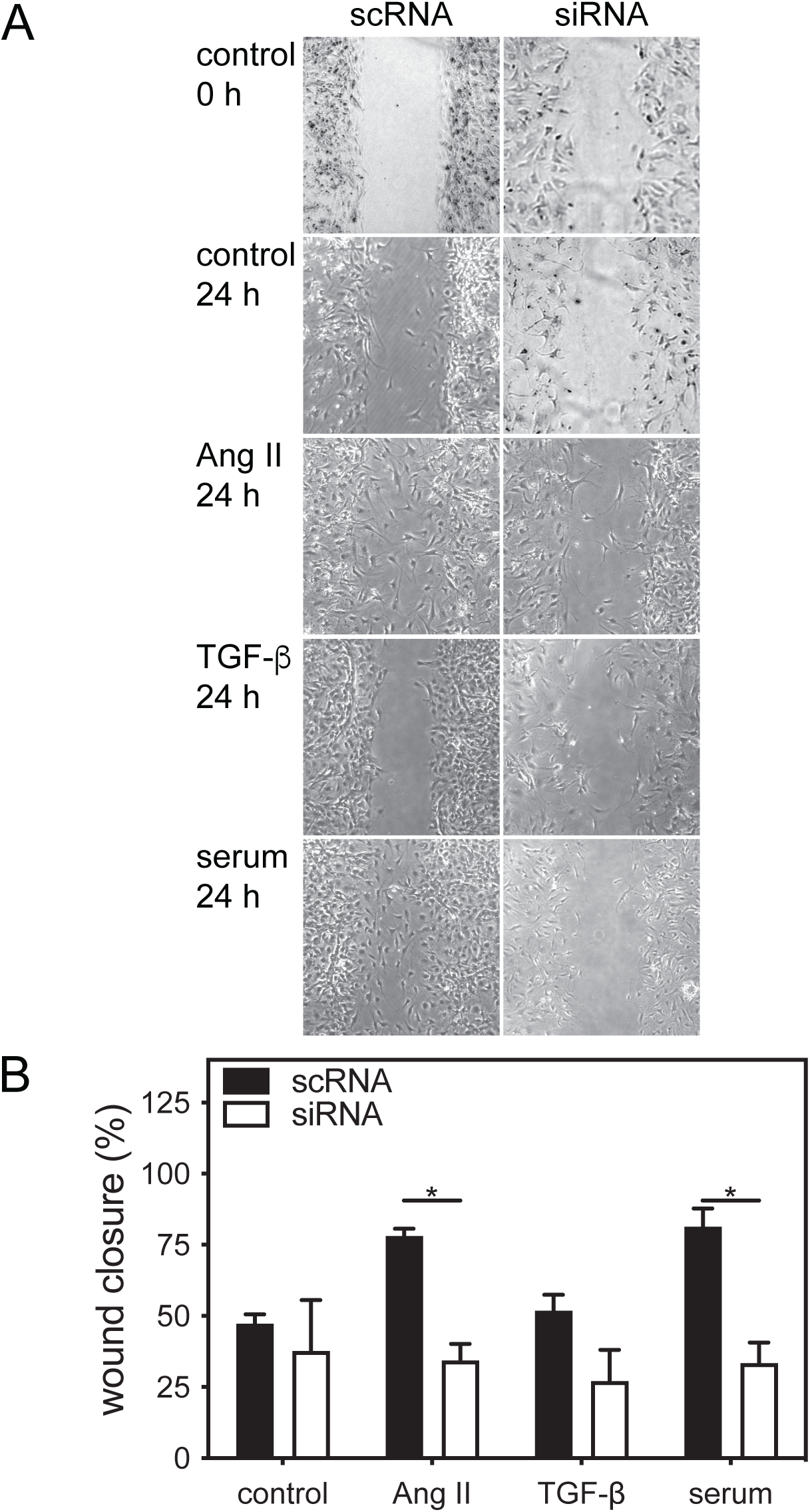
Small interfering (si)RNA-mediated knockdown of ERK3 impairs motility in atrial myofibroblasts. (**A**) Cardiac fibroblasts were isolated from the atria of wild type mice, split into two, and transfected with either siRNA for ERK3 (si) or a scrambled RNA sequence (sc), and motility was assessed by scratch wound assay. After the scratch was imposed, fibroblasts were cultured in serum-free media alone (control) or supplemented with angiotensin II (Ang II; 1 μM), TGF-β (1 μM), or 1% serum. Open wound areas were measured at times 0 and 24 h, and wound closure was calculated. (**B**) Wound closure after 24 h normalized to the mean closure of the control scRNA group (expressed as %). Statistical analysis was by 2-way ANOVA followed Tukey’s post hoc test for multiple comparisons. The data shown are mean ± SEM of assessments performed in fibroblasts isolated from 3 different mice. Bar, 250 μm. \**P* < 0.05.

## 4. Discussion

ERK3/MAPK6 is one of the protein serine/threonine kinases responsible for activating MK5 (16-18, 24), but the physiological role of ERK3 remains poorly understood. The findings of the present study demonstrate that 12-week-old male ERK3 haplodeficient mice have normal systolic and diastolic functions. In response to a chronic increase in afterload, imposed by constriction of the transverse aorta (TAC), cardiac fetal gene expression in ERK3-haplodeficient mice did not differ from wildtype littermates. However, hearts from TAC-ERK3^+/-^ mice showed less hypertrophy than those of TAC-ERK3^+/+^ mice 3 weeks post-surgery. In addition, the increase in *Col1a1* mRNA induced by TAC was attenuated in TAC-ERK3^+/-^ mice compared with TAC- ERK3^+/+^ littermates. Furthermore, as reported previously after shorter-term exposure to increased afterload (19), 3 weeks post-TAC, ERK3, but not ERK4 or p38α, co-immunoprecipitated with MK5 in lysates prepared from both TAC and sham hearts. As 1) ERK3 is an activator of MK5, 2) MK5 immunoreactivity is present in adult cardiac fibroblasts but not myocytes (13), and 3) MK5 mediates cardiac fibroblast function (14, 15), ERK3 may also be involved in regulating aspects of cardiac fibroblast function. Knocking down ERK3 in both quiescent fibroblasts and ‘activated’ myofibroblasts using siRNA prevented TGF-β from increasing the abundance of *Col1a1* mRNA and prevented both Ang II and TGF-β from increasing intracellular and secreted type 1 collagen immunoreactivity in quiescent fibroblasts. Knocking down ERK3 also produced a modest increase in secreted MMP2 activity in quiescent fibroblasts in the presence of Ang II and reduced motility in both ventricular and atrial myofibroblasts. These results suggest a role for ERK3 in regulating fibroblast function and, possibly, reparative and/or interstitial fibrosis.

MK5 haplodeficiency attenuates both hypertrophy and the increase in *Col1a1* evoked by a chronic increase in afterload induced by constriction of the transverse aorta (13) suggesting that some pathological cardiac remodeling involves MK5 activity. The mechanism whereby MK5 is activated has been somewhat controversial and its substrates are largely unknown. There is conflicting evidence about the ability of MK5 and p38 to form a complex *in vivo*. Tandem affinity purification studies did not detect p38α-MK5 complexes in HEK293 cells (25) whereas MK5 and p38 co-immunoprecipitate when over-expressed in NIH3T3 cells (26, 27). Although p38α/β and ERK3/4 have been shown to be upstream of MK5 (16-18, 28, 29), and both ERK4 and p38α immunoreactivity are present in GST-MK5 ‘pull-downs’ from heart lysates (19), ERK3 but not ERK4 or p38α are detected in MK5 immunoprecipitates from mouse heart lysates (19). In addition, 3 days post-TAC, where p38 activity is increased (8), p38α is not detected in MK5 immunoprecipitates whereas ERK3 is (19). Furthermore, in this study, we show that even 3 weeks post-TAC, ERK3 but not p38α or ERK4 form ‘stable’ complexes with MK5. Hence, in the heart, ERK3 and MK5 form, and may function as interacting partners. Interestingly, although ERK3 and MK5 immunoreactivity are detected in heart and fibroblast lysates, MK5 immunoreactivity is not detectable in adult ventricular myocytes (13). Given that both ERK3 and MK5 haplodeficiency results in attenuated hypertrophy in response to TAC, the presence of both ERK3 and MK5 in fibroblasts and the absence of MK5 in myocytes, and the presence of ERK3- MK5 complexes, it is possible that ERK3-MK5 play a role in fibroblast-myocyte signaling during cardiac remodeling that promotes myocyte hypertrophy in response to chronic increases in afterload. This fibroblast-myocyte communication may involve direct cell-cell contact or, alternatively, paracrine signaling.

Chronic increases in cardiac afterload induce cardiac fibroblast proliferation, remodeling of the ECM, and interstitial fibrosis. The major protein constituent of the cardiac ECM is collagen. The fibrillar collagens type I and type III comprise approximately 95% of the cardiac ECM collagens and, of that, type I accounts for approximately 85% (30). Type I collagen is a thick fiber, and it provides high tensile strength and stiffness to the myocardium, whereas collagen type III is thin and flexible, providing more flexibility to the tissue. Although the fibrillar collagens coexist in the cardiac ECM, the ratio of collagen type III to type I determines the functional properties of the heart. Various pathological factors can induce cardiac interstitial fibrosis by enhancing the deposition of collagen or reducing its degradation. Since the primary role of collagen in the cardiac ECM is to maintain the structural framework of the myocardial wall and to facilitate the coordinated mechanical action of the heart, even a slight imbalance in the collagen matrix may have drastic effects on myocardial stiffness and result in decreased compliance (31, 32). In hearts from both MK5 (13) and ERK3 haplodeficient mice, a chronic increase in afterload failed to induce a noticeable increase in collagen accumulation in the interstitium. This was accompanied by a significant reduction in the ability of increased afterload to increase the abundance of *Col1a1* mRNA. Similarly, knocking down ERK3 prevented TGF-β from increasing *Col1a1* mRNA in both quiescent fibroblasts and myofibroblasts. Mechanistically, it remains to be determined if this reflects changes in transcription, mRNA stability, or both. Interestingly. MK5 haplodeficiency did not increase mortality in mice following myocardial infarction, although scar size and collagen content were reduced, suggesting that reparative fibrosis was not impaired (14).

The matrix metalloproteinases (MMPs) are a major class of ECM-degrading endopeptidases that play a significant role in ECM turnover (23, 33-35) and cardiac fibroblasts are a significant source of MMPs. The gelatinases such as MMP-2 and MMP-9 play an essential role in ECM homeostasis by degrading collagen, fibronectin, elastin, and laminin in the myocardium (23, 33, 36, 37). The present study reveals that TGF-β reduced MMP2 activity in quiescent fibroblasts in the presence or absence of ERK3 whereas knocking down ERK3 increased secreted MMP2 activity in these cells in the presence of Ang II. On the other hand, MMP9 activity in cardiac fibroblasts was not affected by ERK3 knockdown. However, the activities of MMPs are tightly regulated by tissue inhibitors of metalloproteinases (TIMPs), endogenous inhibitors of MMPs, and the net proteolytic activity of MMPs is also associated with MMP/TIMP ratio (38, 39). Since cardiac fibroblasts are the cellular source of fibrillar collagen in the heart, and the reduction of type I collagen observed in both ERK3^+/-^ hearts after chronic pressure overload and ERK3-deficient quiescent fibroblasts in response to Ang II and TGF-β stimulation suggest the role for ERK3 in collagen synthesis in cardiac fibroblasts.

The activation of quiescent fibroblasts to myofibroblasts is an important component of myocardial repair following injury. In addition to enhanced production of ECM proteins, including collagens, myofibroblasts display increased proliferation and migration, which are regulated by MAPKs (15, 40). Although the role of ERK3 in cell migration, and the mechanism whereby it exerts its effect, are controversial, ERK3 has been shown to regulate cell migration in multiple cells via kinase-dependent and-independent mechanisms (41-45). Increased ERK3 abundance or activation is associated with increased migration of breast cancer and head and neck cancer cells, whereas ERK3 deletion or knockdown decreases the migration and invasion of lung, cervical, and mammary cancers or cancer cell lines (41, 42, 46-49) yet enhances the migration of myeloma or A-431 cell lines (50, 51). In the present study, siRNA-mediated knockdown of ERK3 reduced the motility of cardiac fibroblasts in response to angiotensin II, TGF-β, or serum without altering basal motility, which is consistent reports of a MK5 deficiency impairing cardiac fibroblast migration (14). These results suggest co-ordinated ERK3-MK5 activation may regulate cardiac fibroblast motility.

TGF-β is a pleiotropic cytokine that is involved in a wide range of cell functions such as cell proliferation, differentiation, migration, inflammation, and ECM deposition (52-55). In mammals, there are 3 TGF-β isoforms: TGF-β1, TGF-β2, and TGF-β3. TGF-β_1_ predominates and is expressed ubiquitously compared to the other isoforms that display more restricted patterns of expression (55-58). Although three distinct genes encode these isoforms, they are structurally related with similar properties *in vitro*, but diverse and context-dependent effects *in vivo* (55-58). Cardiac myocytes and fibroblasts not only express TGF-β and its receptors but are also highly responsive to the TGF-β family members. TGF-β induces cardiomyocyte hypertrophy whereas the fibroblast response is hyperplasia (59-61). However, TGF-β also promotes fibroblast activation to myofibroblasts. The increase in TAC-induced *Tgfb3* mRNA was attenuated in both MK5- (13) and ERK3-haplodeficient mice. Although an elevated and prolonged induction *Tgfb3* mRNA has been reported in experimental models of myocardial infarction (62) and a polymorphism of TGF-β_3_ is associated with changes in LV geometry in hypertensive patients (63), very little is known about its function in cardiac remodeling. Being pleiotropic in its action, TGF-β can elicit both detrimental and beneficial effects under varying pathological conditions, which restrict them from being a target for therapeutic intervention. In the present study, in both quiescent fibroblasts and myofibroblasts, knocking down ERK3 prevented the increase in *Col1a1* mRNA induced by TGF-β whereas it reduced the increase in abundance of intracellular, but not secreted, type 1 collagen immunoreactivity induced by Ang II or TGF-β. Alternatively, knocking down ERK3 increased secreted MMP2 activity in myofibroblasts treated with Ang II but not those treated with TGF-β. In contrast, as mentioned above, the absence of ERK3 reduced the ability of Ang II, TGF-β, or serum to increase fibroblast motility without affecting the basal rate of wound closure. Although the present study used siRNA to knock down ERK3 in fibroblasts, there is considerable evidence for the endogenous post-transcriptional regulation of ERK3. Numerous microRNAs (miRs) target *Mapk6* RNA (miR-26a-5p, −98−5p, −128−3p, −138, −144-3p, −196a, −374a-5p, −495-3p, −499a, −653-5p, −654-3p, −let-7i, −let7f-5p) (48, 49, 64-77). In certain types of cancer or cancer cell lines, miR downregulation results in increased ERK3. In addition, there are miRNA ‘sponges’, such as long non-coding RNA (lncRNA) NEAT1 that binds miR- 495-3p and miR-98-5p (64, 71), circular RNA circ_0000285 that binds miR-654-3p (72), lncRNA SNHG6 that binds miR-26a-5p (67), lncRNA GAS6-AS2 that binds miR-144-3 (66), lncRNA SNHG1 that binds miR-196a (73), and circular RNA circ_0000285 that binds miR-654- 3p (72), which sequester target miRs. Furthermore, the circular DNA circDNAJC11 binds the RNA-binding protein TAF15 to increase the stability of *Mapk6* mRNA (78). Each of these also result in increased ERK3 and this increase is associated with phenotypic changes such as increased proliferation, migration, and/or invasion. miR-mediated posttranscriptional regulation of ERK3 levels has also been demonstrated to play a role in smooth muscle cell proliferation and graft neointimal hyperplasia (69), neuroinflammation and neuropathic pain (77), and ischemia/reperfusion injury (71, 76). In HK-2 epithelial cells, the TGF-β1-induced increase in collagen 1α1, fibronectin, and αSMA mRNA and immunoreactivity are attenuated by an miR- 374a-5p agomir (79). Interestingly, TGF-β1 reduced the abundance of miR-374a-5p. *In vivo*, exosomal-delivered miR-374a-5p agomir inhibits the progression of renal fibrosis (79). However, whereas the miR-374a-5p agomir reduced the abundance of *Mapk6* mRNA in HK-2 cells, in both renal tissue and HK-2 cells phosphorylated ERK3 immunoreactivity was reduced whereas that of total ERK3 was unaffected, indicating an effect on ERK3 activity but not an actual knockdown at the protein level (79). Hence, ERK3 may be involved in multiple aspects of cellular signalling to mediate the effects of extracellular stimuli; however, the role of ERK3 may be cell-type specific and in cardiac fibroblasts the role of ERK3 may also be dependent on the activation state of the fibroblasts.

## 5. Conclusions

The present study highlights the role of ERK3 in cardiac function with specific attention to cardiac fibroblasts. We have shown that although ERK3^+/-^ hearts hypertrophied in response to a chronic increase in afterload, both hypertrophy and the increase in *Col1a1* mRNA were attenuated in hearts from ERK3 haplodeficient mice. We further showed that ERK3 and MK5 form a stable complex in the mouse heart and remain associated even after 3 weeks of increased afterload. This study also revealed the role of ERK3 in fibroblast function, under certain conditions, including collagen production and motility. Further studies are required to provide mechanistic insight into how ERK3-MK5 regulates fibroblast function.

## 6. Limitations of the Study

The ERK3-deficient mice used in this study were a global knockout and haplodeficient mice were used due to neonatal mortality (20). These mice were generated by inserting *LacZ* into exon 2. A more recent study using a conditional knockout where exon 3 of ERK3 was flanked by loxP observed that germline deletion of ERK3 resulted in ERK3-deficient mice that were viable (80). In addition, our previous studies revealed phenotypic differences between MK5 deficient and haplodeficient fibroblasts (15). It would be better to study the role of ERK3 in cardiac remodeling using an inducible, fibroblast-targeted knockout model, as a greater decrease in the abundance of ERK3 could be obtained. The present study examined mice 3-weeks post-TAC. In previous studies, we examined the effects of MK2 deficient 2- and 8-weeks post-TAC and observed that hypertrophy was delayed but not reduced in this model whereas MK5 haplodeficient mice showed no effect on hypertrophy 2-weeks post-TAC but attenuated hypertrophy was observed after 8 weeks (11, 13). Studying the effect of ERK3 deficiency at a longer time point post-TAC would give a better idea of the long-term consequences of ERK3 deficiency on cardiac remodeling. Finally, the studies described herein were undertaken in male mice or fibroblasts isolated from male mice and should be extended to include female mice and fibroblasts isolated from female mice.

## Acknowledgements

We thank Dr. Robert Parent and Adam Sanscartier for animal care and breeding. These studies were supported by Grants-in Aid from the Heart and Stroke Foundation of Canada (G-14- 0006060, G-18-0022227 (BGA) and an operating grant from the CIHR (PJT-173466) to BGA.

## Notes

### Competing Interest Statement

The authors have declared no competing interest.

